# Single Cell Spatial Transcriptomics Reveals Immunotherapy-Driven Bone Marrow Niche Remodeling in AML

**DOI:** 10.1101/2025.01.24.634753

**Authors:** Gege Gui, Molly A. Bingham, Julius R. Herzog, Abigail Wong-Rolle, Laura W. Dillon, Meghali Goswami, Eddie Martin, Jason Reeves, Sean Kim, Arya Bahrami, Hermann Degenhardt, George Zaki, Prajan Divakar, Edward C Schrom, Katherine Calvo, Christopher S. Hourigan, Kasper Hansen, Chen Zhao

## Abstract

Given the successful graft-versus-leukemia cell treatment effect observed with allogeneic hematopoietic stem cell transplant for patients with refractory or relapsed acute myeloid leukemia, immunotherapies have also been investigated in the nontransplant setting. Here, we use a multi-omic approach to investigate spatiotemporal interactions in the bone marrow niche between leukemia cells and immune cells in patients with refractory or relapsed acute myeloid leukemia treated with a combination of the immune checkpoint inhibitor pembrolizumab and hypomethylating agent decitabine. We derived precise segmentation data by extensively training nuclear and membrane cell segmentation models, which enabled accurate transcript assignment and deep learning-feature-based image analysis. To overcome read-depth limitations, we integrated the single-cell RNA sequencing data with single-cell-resolution spatial transcriptomic data from the same sample. Quantifying cell-cell distances between cell edges rather than cell centroids allowed us to conduct a more accurate downstream analysis of the tumor microenvironment, revealing that multiple cell types of interest had global enrichment or local enrichment proximal to leukemia cells after pembrolizumab treatment, which could be associated with their clinical responses. Furthermore, ligand-receptor analysis indicated a potential increase in *TWEAK* signaling between leukemia cells and immune cells after pembrolizumab treatment.

**Highlights:**

- Spatial transcriptomic analysis of R-AML bone marrow niches provides detailed information about intercellular interactions in the tumor microenvironment.
- Immunotherapy shifts the cell composition of the leukemia neighborhood.

**Graphical abstract:** 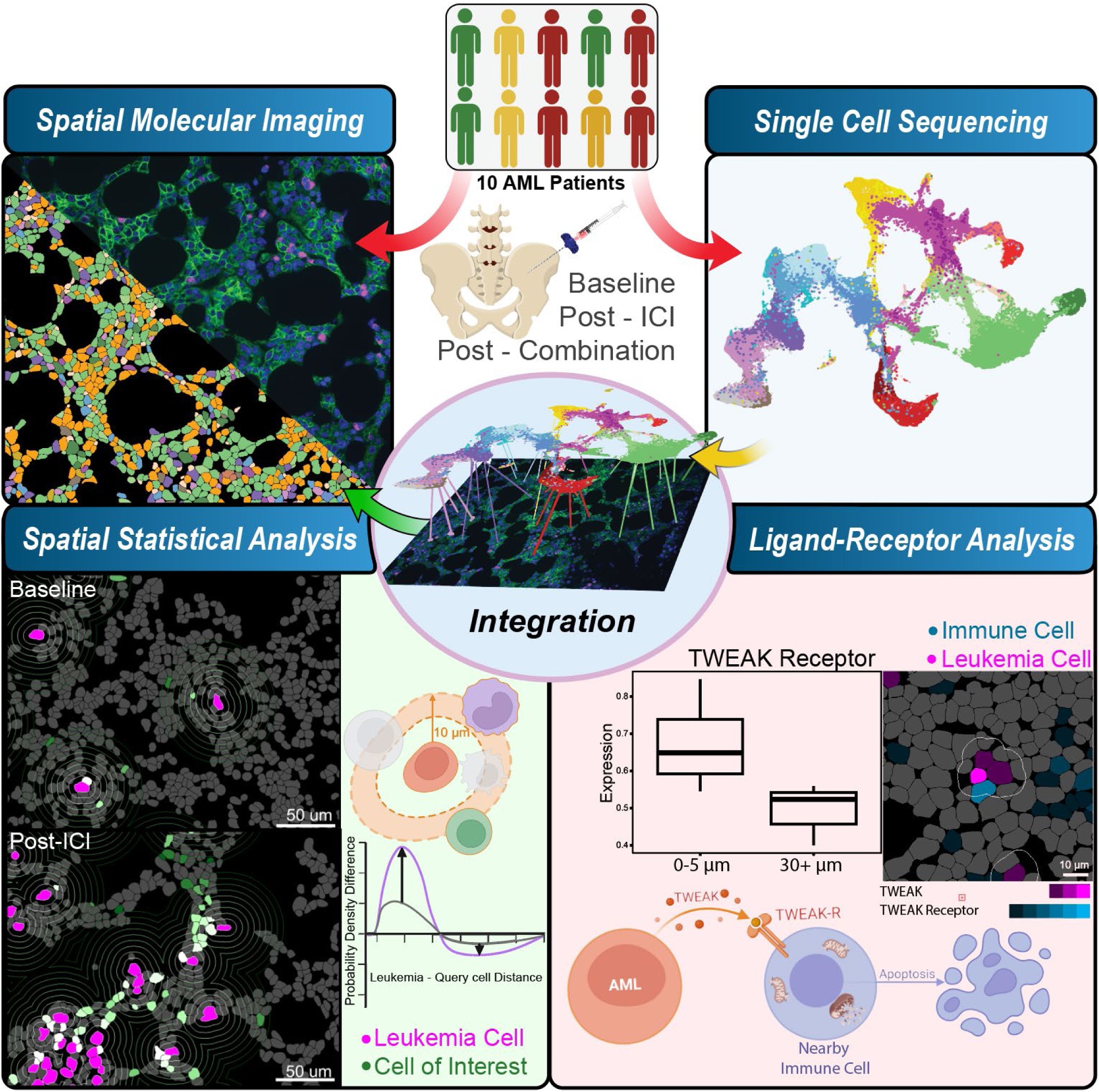

## Introduction

Acute myeloid leukemia (AML) is a deadly hematological malignancy with an estimated 5-year survival rate of 30%^1^. Patients with refractory or relapsed AML (R-AML) have an extremely poor prognosis^2^. Success in using immune checkpoint inhibitors (ICIs) of PD-1 signaling in other cancer types motivated testing of this approach in patients with R-AML^3^. We previously conducted a clinical trial to treat 10 patients with R-AML using the novel combination of pembrolizumab (an ICI) and the hypomethylating agent decitabine (NCT02996474) and reported transcriptomic and proteomic changes of T cells in these patients throughout the treatment course^4,5^. Of the 10 patients treated in this single-institution, single-arm, open-label study, 6 had a response of stable disease, partial, or complete response. When we performed sequencing and transcriptional analysis of T cell receptor β, we found clonal expansion of CD8^+^ effector memory T cells in patients who experienced immune-related adverse events^4^. These expanded T cell populations were largely PD-1 positive and expressed transcriptional profiles consistent with T cells in an activated, cytotoxic state. The T cell populations in the two patients with complete or partial responses, however, did not exhibit a distinct transcriptional profile. These prior results, which characterized the transcriptional and immunophenotypic changes in the T cell population but failed to explain the difference in responses by looking at a single cell population, underscore the need to characterize not only individual cell populations but also their dynamic interactions and spatial distribution within the tumor microenvironment.

Studies quantifying immune cell infiltration in tumors have demonstrated its predictive power, showing that patients with a high level of immune cell infiltration in the tumor (also called a “hot” tumor) have superior responses to immunotherapy^6,7^. In addition, spatial signatures and patterns derived from multiplex immunofluorescent antibody staining can predict response in multiple solid tumor types^8–11^. To better understand the complex interactions between tumor and immune cells in R-AML, patients receiving ICIs and hypomethylating agents, comprehensive profiling of different cell types in their spatial context is required. We have used single-cell RNA sequencing (scRNA-seq) in combination with antibody-derived tags (ADTs) to obtain transcriptomic and immunophenotypic information, but this approach lacks the spatial information of the bone marrow niche because cells are dissociated from their tissue context for sequencing analysis. Although traditional hematoxylin and eosin staining, immunohistochemistry, and immunofluorescence imaging preserve the tissue’s spatial context and enable general spatial analysis of leukemia and immune cells, these methods do not provide sufficient information to identify and characterize the diverse immune subpopulations crucial for a comprehensive analysis of the tumor microenvironment^12,13,14^.

Previous attempts to combine RNA detection and protein imaging have proven difficult, as the tissue digestion process required to expose RNA binding regions tends to damage the protein epitopes to which the antibodies bind. Recently, however, as many markers of cellular function that cannot be stained by antibodies are now easily detectable via spatial transcriptomic analysis, its combination with immunofluorescence imaging has allowed us to detect novel cell- cell interactions in the tumor microenvironment.^15^ This enables a more robust understanding of the tumor microenvironment’s structure and heterogeneity, including tumor-immune interactions, and allows us to even further characterize immune cells’ functional states (for instance, we can now identify T cells expressing genes indicative of cytotoxicity or exhaustion). The CosMx Spatial Molecular Imager (CosMx SMI; NanoString Technologies) can image a formalin-fixed paraffin-embedded slide, from which users select fields of view (FOVs) for high-resolution imaging and transcript detection.

Our present work combines traditional scRNA-seq plus ADTs with advanced spatial transcriptomics methods to investigate the spatiotemporal interactions between different immune cell populations and leukemia cells in bone marrow samples collected from six patients with CD34^+^ leukemia before and after immunotherapeutic treatment.

## Results

### Single-Cell Transcriptomic Analysis of Bone Marrow Samples Identifies Cell Type Markers and Distinct Patient-Specific Leukemia Populations

Patients were categorized into two groups according to their clinical response to therapy: responders, which included patients who achieved clinical benefit (complete/partial response or stable disease), and nonresponders, i.e., those with progressive disease (**Figure 1A, Table S1**). Previously collected bulk RNA and protein sequencing data from bone marrow and blood did not identify any significant changes in gene or protein expression across study time points or patient response (**Figure S1**). We therefore conducted further experiments utilizing the previously described advanced tissue characterization technologies to test our hypothesis that immunotherapy remodels bone marrow niche in AML.

**Figure 1:**
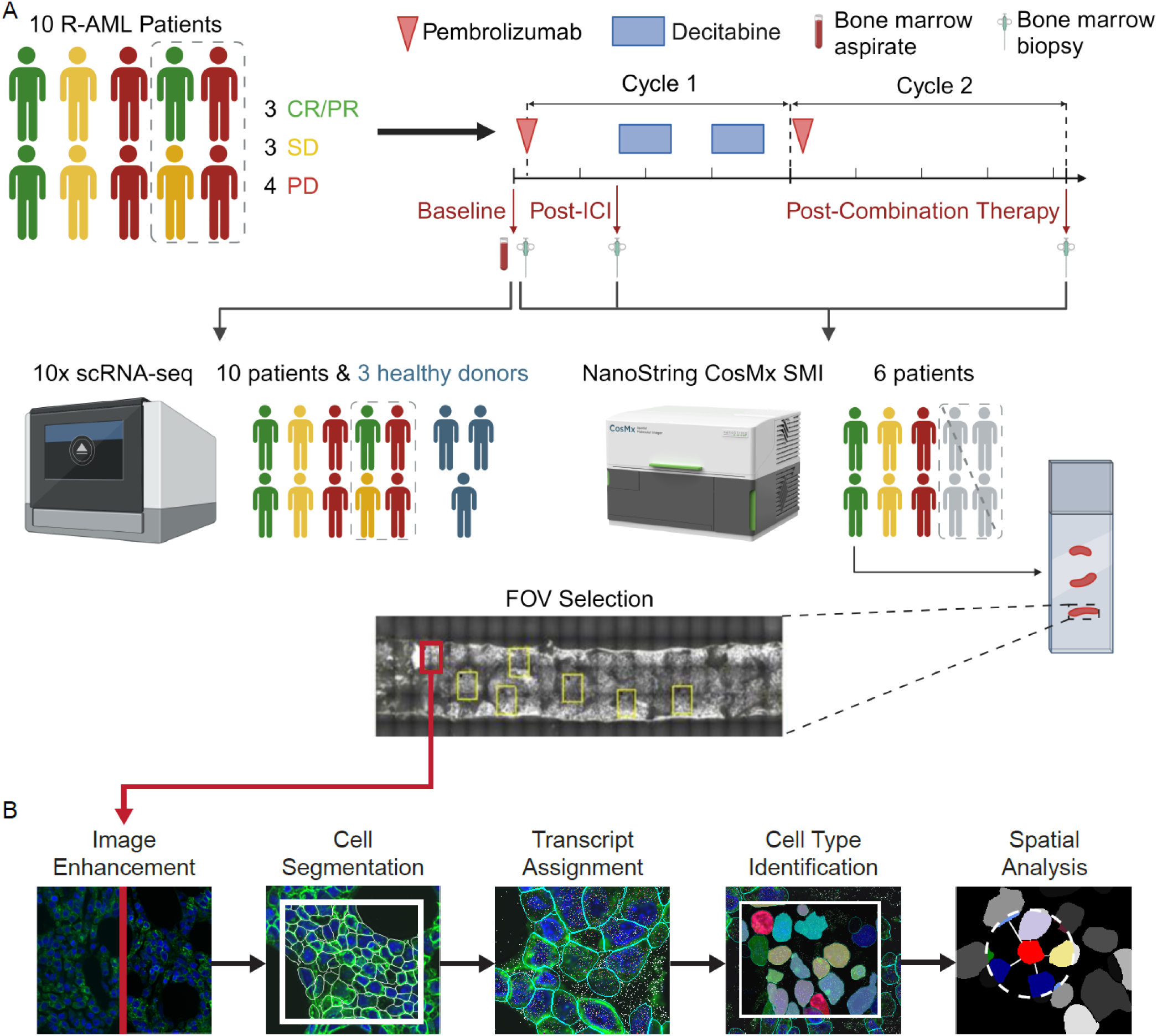
Study scheme and analysis outline. **(A)** Schematic of the study design. Ten patients with R-AML underwent a pretreatment bone marrow biopsy (baseline) and aspirate collection. They then received the ICI pembrolizumab on day 1 of the first treatment cycle. On day 8 of cycle 1, another bone marrow biopsy was obtained (post-ICI time point), after which they started receiving the hypomethylating agent decitabine, which they continued to receive for 5 days. On day 15 of cycle 1, patients began another 5-day decitabine treatment. On day 1 of cycle 2, patients again received pembrolizumab but no other treatment for the remainder of cycle 2. On the last day of cycle 2 (day 21), a third and final bone marrow biopsy was taken from each patient (post-combination therapy time point). The treatment continued for 8 total cycles. 10x scRNA- seq was performed on baseline bone marrow aspirate samples from all 10 patients. scRNA-seq results were used to create cell-type-specific gene expression profiles for each patient. Of the 10 patients, biopsies from 6 with CD34^+^ leukemia were used for the spatial transcriptomic assay. Each patient’s three biopsies were loaded onto a single slide and 5 to 15 FOVs were selected for analysis on the CosMx SMI. **(B)** The image processing pipeline used for analysis consisted of image enhancement, cell segmentation, assignment of transcripts to individual cells, cell type identification, and further spatial analyses. The cell-type-specific gene expression profiles generated for each patient from the 10x scRNA-seq data were used to identify cell types in the SMI data.

We obtained baseline bone marrow aspirate samples from all 10 patients in the clinical trial for 10x Genomics 3’ scRNA-seq with 77 ADTs. Three bone marrow samples from healthy donors were used as the reference for data integration and cell type annotation (**Figure 1B**). Each sample underwent individual quality control and normalization, whereupon their data was integrated, resulting in 55,635 total cells (**Figure 2A**, **Figure S2A, Table S3**).

**Figure 2:**
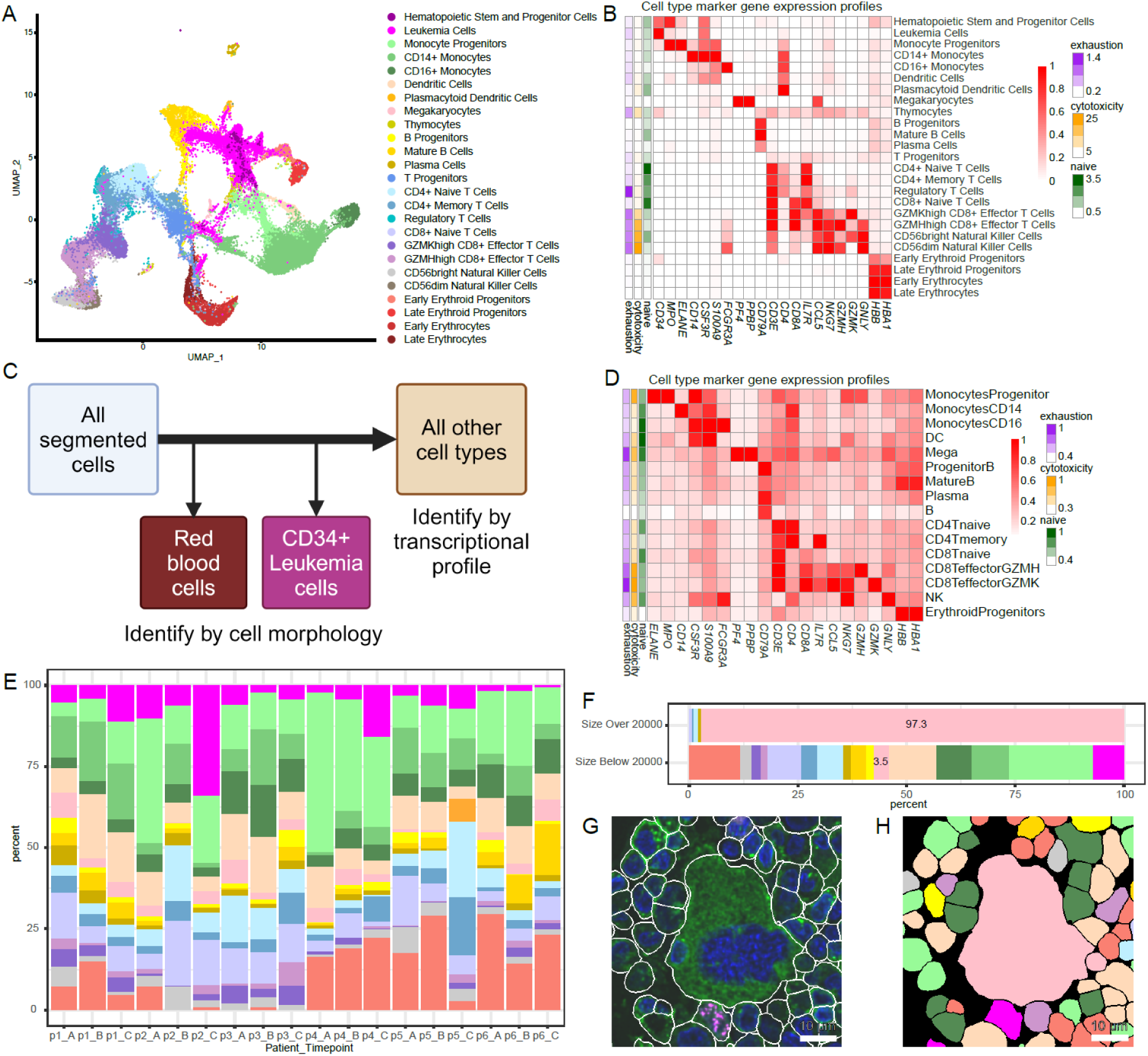
Cell type annotation and visualization for 10x single-cell RNA-seq with cell surface antibodies and CosMx platforms. **(A)** UMAP visualization of all cells from 10 patient and 3 healthy donor samples by 10x with cell type annotation. **(B)** Normalized marker gene expression across different cell types by 10x. **(C)** Cell typing workflow for CosMx data. RBCs were removed from further analysis and leukemia cells were identified by CD34 staining. All remaining cells were identified by transcriptional profile. **(D)** Normalized marker gene expression across different cell types by CosMx. **(E)** Cell type distribution across all samples for 6 patients at 3 time points of CosMx data. **(F)** Comparison of cell type percentages for cells larger or smaller than 20,000 pixels in area. Those annotated as megakaryocytes have relatively larger cell size. **(G)** A megakaryocyte identified by size. Scale bar represents 10 microns. **(H)** Cell typing result in the same cropped region in which megakaryocytes are shown in light pink. Scale bar represents 10 microns.

The median number of cells for all samples was 4,579 (range: 2,517-6,207), with a median number of transcripts of 3,787 (IQR: 2,335-5,997) and a median number of ADTs of 599 (IQR: 390-904). Unsupervised clustering of the integrated data did not show sample-specific or response-specific patterns, mainly separating healthy donor cells from patients (**Figure S2B**). Differential protein expression between clusters consisting only of cells from either healthy donors or patients confirmed that the cells in the cluster dominated by patient cells had enriched stem cell-related protein signatures, such as CD34 and CD117 (**Figure S2C**). Meanwhile, the cluster of predominantly healthy donor cells was annotated as consisting of mature B cells, a population typically diminished in patients with AML (**Figure S2D**).^16^ Cell types of all clusters were annotated by using the top differentially expressed ADT signatures, which were further supported by the top gene markers (**Figure 2B**). These gene markers were used to identify reference cells in the downstream spatial transcriptomic analyses. A total of 5,468 cells were labeled as leukemia cells, consisting of 13.8% of all patient cells (range per patient: 3%-38%) after excluding erythrocytes.

The clinical data suggested that patients had different leukemia-associated immunophenotypes (LAIPs) and cytogenetics at initial diagnosis^4^. These differences would be useful for validation of our malignant cell identification. The data from each individual patient was subset and separately underwent dimension reduction, clustering, and cell type annotation, showing that the LAIP of patient-specific malignant cell clusters matched what was reported clinically (**Figure S2E**)^4^. Additionally, 91% of the leukemia cells identified in the integrated dataset overlapped with the patient-specific malignant cells, demonstrating agreement between the two assignment methods. Most of the leukemia cells identified in individual samples but not the integrated dataset were progenitor cells: 78% were early erythroid precursors, monocyte progenitors, thymocytes, and T cell progenitors. Patient 3 and patient 6 were selected for copy number variation analysis because their cytogenetic analyses discovered chromosomal loss or gain in 20 metaphases. Compared to gene expression from hematopoietic stem and progenitor cells from healthy donors, the posterior estimation of copy number changes in the malignant cells from patient 3 showed deletions in chr5/chr7 and gain in chr21, consistent with the cytogenetic report at diagnosis (**Figure S2F**). Although the population of leukemia cells was smaller in patient 6, the results still matched their clinical reports and showed a loss in chr7 and gain in chr8.

### Further Training of Combined Nucleus-Cell Membrane Model Yields Accurate Cell Segmentation

Before cell type classification and spatial analysis of the tumor microenvironment with CosMx data is possible, the detected RNA transcripts must be assigned to single cells, which must first be detected via cell segmentation. NanoString uses Cellpose^17,18^, an open-source, state-of-the- art segmentation algorithm for cell microscopy data, which is meant to detect all the cells and identify their boundaries in each FOV. Cellpose is among the most accurate cell segmentation algorithms available, and yet the NanoString-generated segmentation of our CosMx SMI data did not always accurately represent cell borders. Comparing their segmentation to manually annotated ground truth showed that the annotated borders were significantly expanded to capture more transcripts than were truly inside the cell. Additionally, cells in our dataset exhibiting unique morphology, such as endothelial cells and megakaryocytes, were often not accurately segmented.

To improve the segmentation accuracy and to fully capture the diversity of cell types in our imaging data, we trained our own nuclear and membrane segmentation models using the Cellpose 2.0 “train-your-own model” feature^17^. We generated a large training dataset by manually annotating 448,439 nuclei and 318,401 cell membranes based on both the nuclear and cell membrane staining of the CosMx SMI data, using these comprehensive datasets as the ground truth for training and eventually testing our model (**Figure S3A-F**).

Additionally, we created an algorithm to combine nuclear and membrane segmentation results such that each cell could be represented by a single mask with a single-cell identification. It was important to train both nuclear and membrane segmentation models because while membrane segmentation most accurately represents a cell’s true border, some cells had very weak or no visible membrane staining, which were likely due to staining artifacts. In these cases, detected nuclei were expanded to estimate the cell’s border (**Figure S3G**).

To evaluate the performance of our segmentation and merging process, we compared its F1 score to that of the default segmentation generated by NanoString from the CosMx data. The F1 score, which balances precision and recall, was calculated at an intersection-over-union (IOU) threshold of 0.7 (see Methods for detailed calculation). The optimal IOU threshold to use for model evaluation depends on the application. Typical IOU thresholds range from 0.5 to 0.9, but 0.7 is a common IOU used to evaluate cell segmentation methods^19^. A 0.5 IOU was too lenient for our purposes, as it only required a true-positive segmentation mask to overlap 50% with the actual cell. On the other hand, 0.9 was too stringent, calling detected objects with less than a 90% overlap between mask and true cell false positives. At an IOU threshold of 0.7, our segmentation result showed significant improvement in accuracy compared to the original CosMx SMI platform output^15^ in our images: the F1 score of the NanoString segmentation was 0.23, while our new approach attained an F1 score of 0.57 (**Figure S3H**).

Our trained and merged segmentation result not only improved the F1 score but also better captured the variability in cell size present in our dataset. This was a key area for improvement, as our images contained a diversity of cell types with vastly different morphologies, ranging from small, round lymphocytes to large, multilobed megakaryocytes. NanoString’s default segmentation result detected a total of 593,279 cells across all six patients and our trained nuclear-membrane model combination detected 629,051 cells (6% more than NanoString’s default segmentation result). Furthermore, cells segmented by our method were on average smaller than those in NanoString’s segmentation, which generated segmentations with a median size of 58.3 µm^2^ (range: 54.2 µm^2^ ∼ 70.1 µm^2^) while our method identified cells with a median size of 46.1 µm^2^ (range: 41.6 µm^2^ ∼ 57.0 µm^2^). Since we segmented nuclear and membrane image channels separately, we were able to limit our use of nuclear expansion to cells without a detectable membrane, leading to fewer instances of cell size inflation via over-expansion, which improved the accuracy of transcript-based cell characterization (**Figure S4**).

### Cell Typing *in situ* by Protein Expression, Transcriptomic Profile, and Morphological Features

Before cell typing the CosMx data, we used several criteria to assess the heterogeneity of the samples. The transcripts and CD34 protein channel were plotted by the measured local coordinates per FOV, which identified different cell densities and distributions across FOVs, showing the heterogeneity of cell distributions within the bone marrow biopsy and background tissue binding (**Figure S5A**). There were 141 FOVs in total (range, 21-25), and a median of 4,146 cells per FOV (range, 1,041-9,408). The average transcripts per cell showed strong sample- and FOV-specific variations (**Figure S5B**), with a median of 102 considering all FOVs (IQR: 45-177).

Data from the CosMx SMI provides the unique opportunity to capture both the morphology and transcriptome of a given cell, enabling more robust cell type assignment and validation than would be possible from single-cell transcripts alone. Before proceeding with transcript-based cell type identification, we had to identify red blood cells (RBCs) by morphology and remove them from the downstream analysis (**Figure 2C, Figure S6**). Even though RBCs lack a nucleus and are not clearly stained by the B2M/CD298 membrane marker, their autofluorescence makes them visually detectable. After segmentation, we fed cropped images of each individual cell’s B2M/CD298 and DAPI channels into EfficientNet^20^, a convolutional neural network (CNN) designed to classify images. As its final output, EfficientNet provides a vector of probabilities, known as an embedding, that quantifies how likely the input image is to belong to one of 1000 predefined classes. Rather than letting the classifier run to completion, we extracted the 672 features used to describe each single-cell image from the second to last (B0) layer of the CNN. We then used Phenograph^21^ for unsupervised clustering to cluster cells by their extracted features, independently for samples from each patient^22^. By labeling each cell in the image with its cluster number, we were able to identify which clusters were predominantly composed of RBCs. The number of RBCs varied per patient: 5,315 RBCs were identified among segmented cells in FOVs from patient 1, 26,881 from patient 2, 4,624 from patient 3, 1,476 from patient 4, 30,850 from patient 5, and 1,437 from patient 6. After the RBCs were identified by morphology, they were removed from subsequent analysis. Most of the morphologically identified RBCs had low gene expression, with 95% having a total gene count of 44 or fewer.

Since all selected patients had CD34-positive leukemia cells, we used the CD34 protein channel information to identify leukemia cells. We once again used the EfficientNet classifier to extract embeddings from cropped images of every cell in the dataset, but rather than including the B2M/CD298 and DAPI channels, the crops included only the CD34 protein channel of each segmented cell. Leukemia cells were thereby identified based on cell morphology and the CD34 protein expression pattern, which enabled us to exclude CD34-expressing endothelial cells. Since the clinical information and scRNA-seq data indicated that normal hematopoietic stem cells were extremely rare in these samples, by this method a total of 40,112 leukemia cells were identified via unsupervised clustering: 5,005 from patient 1, 19,833 from patient 2, 5,005 from patient 3, 6,036 from patient 4, 2,332 from patient 5, and 1,901 from patient 6 (**Figure S7**).

After the annotated RBC and leukemia cell populations were removed, additional low-quality FOVs and cells were removed from the analyses and the genes in the spatial gene panel which had been previously identified as cell type markers in the 10x scRNA-seq data were used to select reference cells. These overlapping genes included *HBB* and *HBA1* for erythroid progenitors; *PF4* and *PPBP* for megakaryocytes; *IL3RA*, *CD33*, and *LYZ* for dendritic cells (DCs); *CD3E*, *CD3D*, and *CD3G* for T cells; *CD19* and *CD79A* for B cells; *MPO* and *ELANE* for monocyte progenitors; *CD14*, *CSF3R*, *S100A9*, *CD33*, and *LYZ* for CD14^+^ monocytes; *GNLY*, *NKG7*, and *FCGR3A* for natural killer cells; and *FCGR3A* (without *GNLY* or *NKG7*) for CD16^+^ monocytes. The lymphocyte category was further subdivided based on the expression of *CD4*, *CD8A*, *CD8B*, *IL7R*, *CCR7*, *CCL5*, *GZMH*, and *GZMK* for T cells and *MS4A1*, *TCL1A*, and *JCHAIN* for B cells. Cells expressing these overlapping genes most highly were selected as the reference cells for the cell type prediction algorithm InSituType^23^, which annotated cell types for the remaining cells. By using the posterior probability of the InSituType output, cells were annotated with the expected differential marker gene expression (**Figure 2D-E**). To validate the inferred cell type annotation, we leveraged imaging data-extracted cell size information and the presence of large megakaryocytes, finding that a total of 98% of cells larger than 648 µm^2^ (20,000 pixels) were correctly labeled as megakaryocytes (**Figure 2F-H**).

### Representing Leukemia Cell Neighborhoods with Linear Mixed Model Reveals Shifts in Cell Composition After ICI Treatment

Common methods of studying cell-cell and neighborhood interactions within the tumor microenvironment begin by computing the distance between each cell and all other cells. Currently, it is a common practice to reduce each cell to a single coordinate location by finding its centroid, and then measuring the distance between each cell’s centroid and all other cell centroids^24^. However, this method has several limitations: not only are irregular cells not accurately represented by a simple centroid location, but centroid measurements are also unable to accurately quantify the number of directly touching cells (**Figure S8**). When examining the composition of each leukemia cell’s neighborhood, we developed a new computational approach to quantify cell-to-cell distances using the distance between the closest points of each cell’s edge, as opposed simply the distance between centroids. After applying this method to our dataset to count the number of cells in direct contact with each cell, we found that the mode of this measurement was three directly touching cells, which represents three equidistant neighbors. Had we used centroid-to-centroid distances, these three neighbors would have been ranked with respect to their proximity to the reference cell, which would be an inaccurate representation of the biological reality.

The spatial transcriptomic data enabled us to determine whether there were any changes in cell type abundance in the regions surrounding leukemia cells specifically. To study the cell type density changes around leukemia cells across patient response and study time points, we fit a series of Poisson linear mixed effect models to the count data extracted for each cell type with patients grouped as responders and nonresponders. This approach accounts for the count nature of our data while considering the repeated measures structure and potential variability between patients. In doing so, we considered that patients who responded similarly to the ICI would display similar cell distribution patterns with surrounding leukemia cells. Ring-shaped neighborhoods at radial distances of 0-5, 5-15, 15-25, 25-35, and 35-45 µm were defined around the cell membrane of each leukemia cell (**Figure 3A**), and the abundance of each different cell type in each of these radial neighborhoods analyzed as the dependent variable in the linear mixed model. We used the total number of cells inside each neighborhood as the offset. This enabled us to compare the prevalence of different cell types at different proximities to a leukemia cell, and to determine whether their proportions varied across study time points and/or with increasing distance from the central leukemia cell.

**Figure 3:**
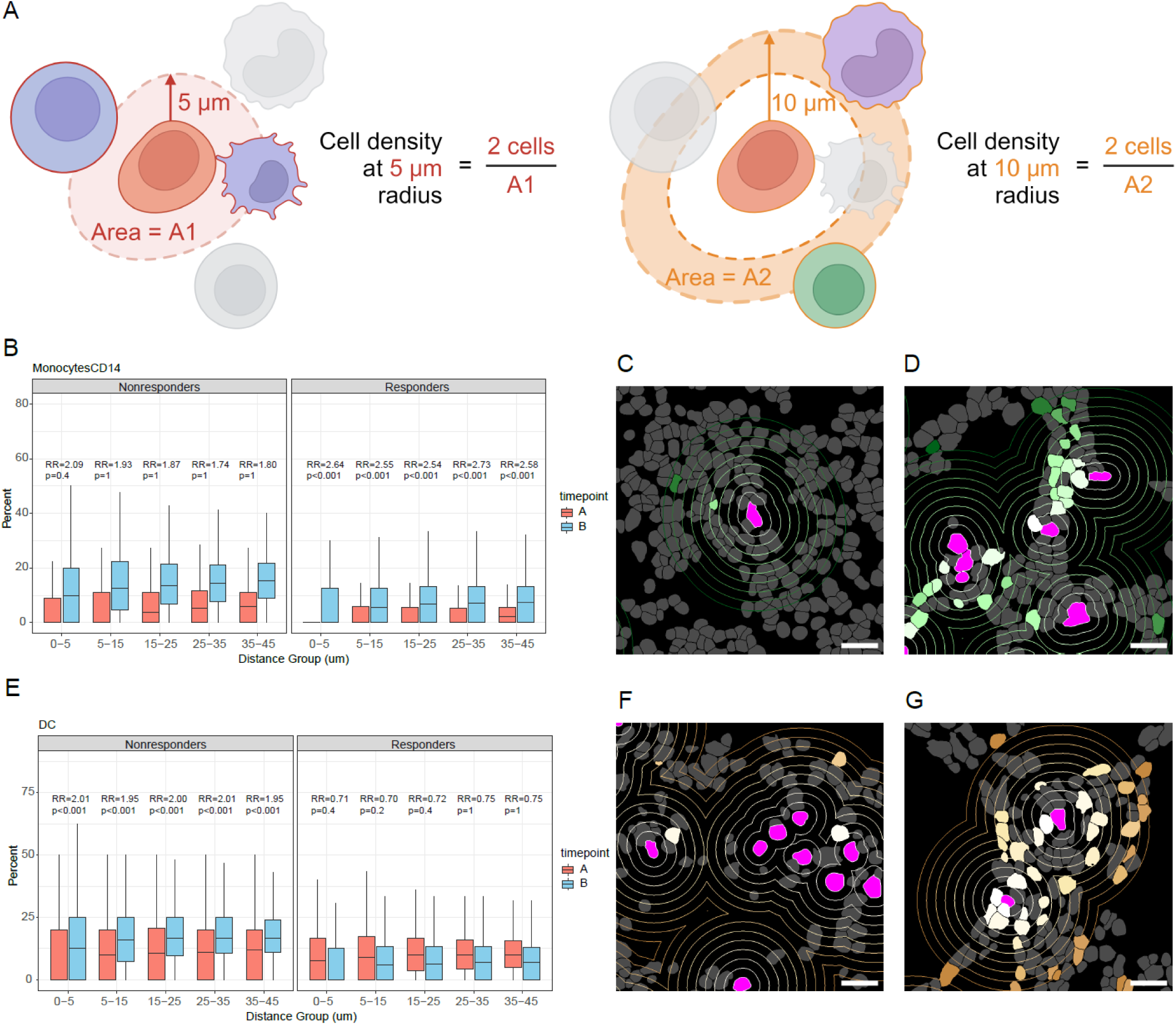
Comparison of leukemia cell neighborhood composition by time point and response group. **(A)** Schematic depicting how cell type composition was determined in leukemia cell neighborhoods of varying radii. The numbers of cells of each cell type were quantified in neighborhoods 0-5, 5-15, 15-25, 25-35, and 35-45 microns from the edge of each leukemia cell. **(B)** Comparing the densities of CD14^+^ monocytes at increasing distance from leukemia cells between baseline and post-ICI samples among nonresponders (left) and responders (right). In neighborhoods at radii 5-15, 15-25, 25-35, and 35-45 microns from leukemia cells, there were higher proportions of CD14^+^ monocytes in the post-ICI samples, but these trends were significant only in responders. **(C)** Representative image of segmented cells showing CD14^+^ monocyte density around leukemia cells in one responder at baseline. Concentric rings around leukemia cells are placed at increasing distances from each leukemia cell, going from 0-45 microns in 5-micron increments. Leukemia cells are in magenta, whereas CD14^+^ monocytes range from white to light green to dark green as they increase in distance from the nearest leukemia cell. Scale bar represents 20 microns. **(D)** Representative image of segmented cells showing CD14^+^ monocyte density around leukemia cells in one responder in the post-ICI sample. **(E)** Comparison of the densities of DCs among nonresponders (left) and responders (right) at increasing distance from leukemia cells between the baseline and post-ICI samples. Proportions of DCs were higher in the post-ICI samples at radii 5-15, 15-25, 25-35, and 35- 45 microns from leukemia cells in nonresponders. **(F)** Representative image of segmented cells showing DC density around leukemia cells in one nonresponder at baseline. Leukemia cells are in magenta, whereas DCs range from white to light brown to darker brown as they increase in distance from the nearest leukemia cell. Scale bar represents 20 microns. **(G)** Representative image of segmented cells showing DC density around leukemia cells in one nonresponder in the post-ICI sample. RR: rate ratio. All p-values in the figure were adjusted p-values

From the separate models we fit for different cell types considering the interactions between patient response, distance groups, and treatment time point, we were able to investigate how ICI treatment perturbed the prevalence of different cell types within leukemia cell neighborhoods in both responders and nonresponders (**Table S2**). We started with examining whether the proportion of cells of any cell type changes around the nearest leukemia cell when comparing responders with nonresponders at baseline. We observed that there were fewer granzyme K^+^ CD8^+^ T effector cells and CD14^+^ monocytes in leukemia cell neighborhoods in responders than in nonresponders at baseline (**Figure S9A-B**). Previous studies have found that patients with higher levels of classic monocytes before ICI treatment tend to have lower overall survival^27^, which aligns with our observation that CD14^+^ monocytes were more prevalent at baseline among nonresponders. We also observed increased proportions of monocyte progenitor cells in responders than in nonresponders at baseline (**Figure S9C**). Higher numbers of CD14^+^ monocytes were seen at the post-ICI time point than at baseline in responders, suggesting that the elevated numbers of monocyte progenitor cells among responders at baseline may have differentiated into CD14^+^ monocytes after treatment. Studies in mice have found that anti-PD1 promoting can catalyze myeloid progenitors to differentiate into anti-tumor effector cells^28^.

Among responders, the proportion of CD14^+^ monocytes in leukemia cell increased after ICI treatment across all defined radial neighborhoods (rate ratio [RR] range 2.5-2.7, adjusted p- values < 0.001). This trend was observed but not significant in nonresponders (**Figure 3B-D**). This suggested that there was an overall increase in CD14^+^ monocytes post ICI therapy for responders, irrespective to leukemia cell proximity. Classic monocytes express chemokine receptors and are known to migrate toward inflammation^25^. Their generalized influx after ICI therapy in responders suggests a response to inflammation around the leukemia cells.

Among nonresponders, we observed an increase in the proportion of dendritic cells (DCs) in all neighborhoods, suggesting an increase in population at the post-ICI time point compared with baseline (RR range: 1.95-2.01, adjusted p-values < 0.001). The trend was the opposite in responders but was not statistically significant (**Figure3E-G**). It has been established that DCs are of limited utility after activating neighboring T cells by antigen presentation, and in fact their accumulation may yield overactivation of the immune system. Previous studies have shown that DCs can undergo attack and elimination by CD8 T cells after serving as antigen-presenting cells. It is possible that this post-ICI DC accumulation among nonresponders is related to an ineffective CD8 T cell response^26^.

All the findings from these models indicated that the significant differences identified were across all pre-specified distance groups, implying the potential shift in cell distribution related to therapy and clinical response.

### Changes in Patient-Specific Spatial Cell Distribution Are Identified by Density Comparison of Time Points Across Cell Types

The linear mixed model assumed there were consistent changes across patients who responded to the therapy compared with those who did not and used a simple discretization of the cell distances. It also did not account for overlapping neighborhoods. Grouping the patients by their response showed us the high-level trend of cell enrichment around leukemia cells. In the meantime, we observed high heterogeneity across samples **(Figure S3)**; thus, we hypothesized that there could be patient-specific signatures that were obscured by considering the patients in groups. We developed a new algorithm that was able to detect patient-specific shifts in cell type density around leukemia cells between two time points of interest, which we took to be baseline and the end of cycle 2 of treatment (**Figure 4A**, **Figure S10**).

**Figure 4:**
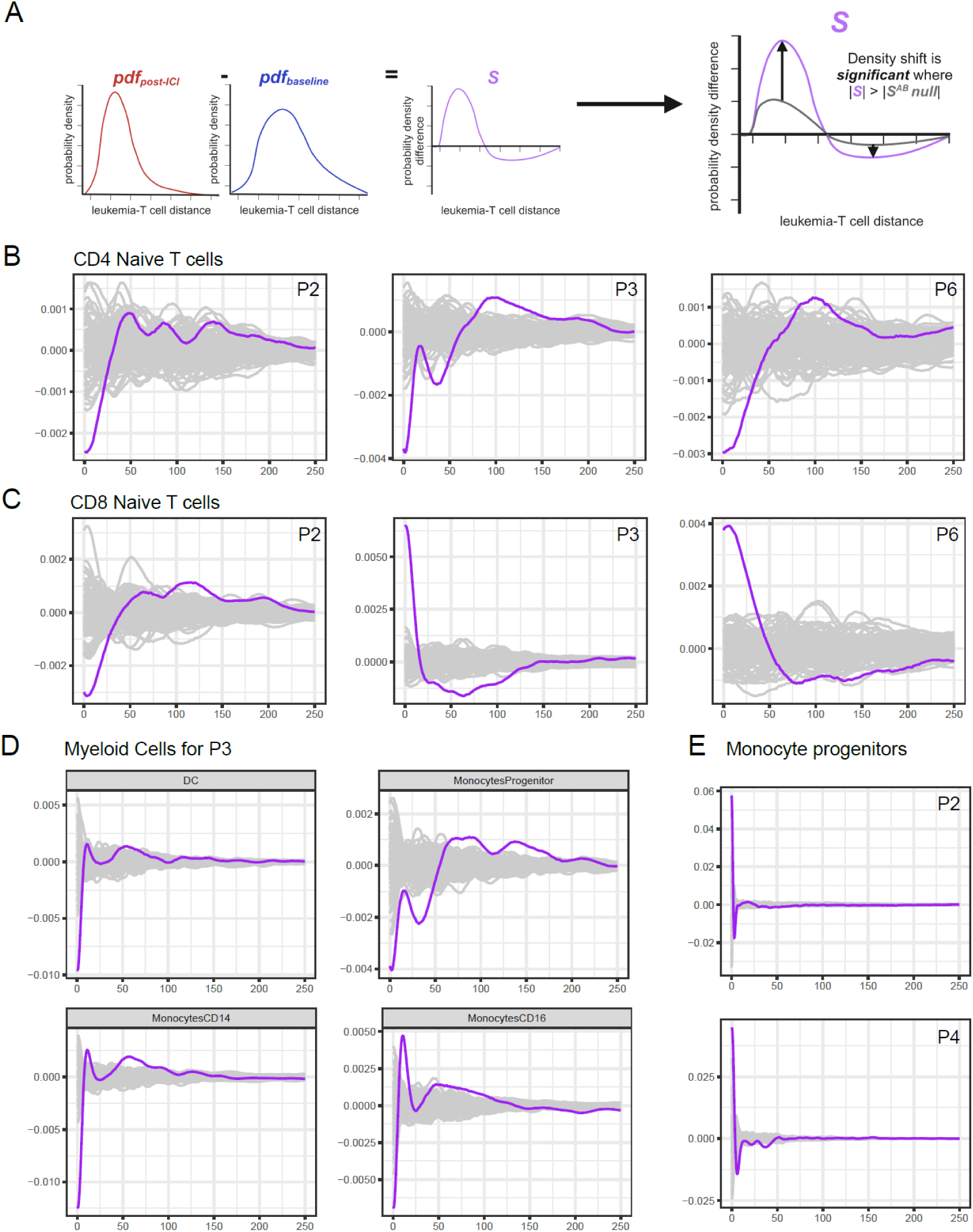
Leukemia-T cell density shift from baseline to the post-ICI time point (purple) compared with simulated background distribution (grey) in pixels. **(A)** Schematic representing this comparison. (B) Density shift in CD4 naïve T cells, **(C)** CD8 naïve T cells, **(D)** myeloid cells for P3, and **(E)** monocyte progenitors. The distance was limited to 45 microns (1 pixel = 0.18 micron) with the density calculated. The plot was normalized to the median of the simulated curves at each distance.

Responder patients 2 and 6 and nonresponder patient 3 exhibited a consistently significant trend of enrichment in the naïve CD4^+^ T cell population at baseline around leukemia cells compared with the end of the second treatment cycle (**Figure 4B**). For the naïve CD8^+^ T cell population, patient 2 showed a trend of having more naïve CD8^+^ T cells at baseline than at the end of cycle 2; however, patients 3 and 6, who had different responses, consistently displayed an opposing trend compared with patient 2, with a positive density shift for CD8-naïve T cells around leukemia cells at the end of cycle 2 from baseline (**Figure 4C**).

There were additional significant shifts in the myeloid cell population density in nonresponder patient 3. There was an enrichment of DCs, monocyte progenitors, CD14 monocytes, and CD16 monocytes around leukemia cells at baseline compared with the end of cycle 2 (**Figure 4D**). For other patients in terms of myeloid cells, patient 2 and patient 4, who were both responders, had a positive density shift in monocyte progenitors with treatment (**Figure 4E**). The other positive shift for patient 3 comparing the two times was for megakaryocytes (**Figure S11**). Patient 6, who was one of the responders, had an increase in progenitor B cells and a decrease in mature B cells **(Figure S11)**.

### Ligand-Receptor Analysis Reveals Leukemia to Myeloid TWEAK Signaling Post-ICI Therapy

Beyond simply understanding the changes in the spatial distribution of cells in the tumor microenvironment, we wanted to identify ligand-receptor pairs involved in the interactions between leukemia cells and other cell types nearby. Given the relatively low read counts of our dataset, especially compared with traditional scRNA-seq studies, we utilized a pseudo-bulk approach. We aggregated transcripts captured within leukemia cells of each FOV to determine which ligand or receptor genes were more highly expressed in leukemia cells than in other cell types. Then for each individual cell type, we aggregated transcript counts from all cells of that type located within 5 microns of the nearest leukemia cell. To determine whether the complementary ligand/receptor gene to that expressed by the leukemia cell was enriched in leukemia cell neighbors specifically and not just in all cells of that type across the FOV, we compared ligand/receptor gene expression in this group of cells within 5 microns of a leukemia cell with the expression of the same transcript in all cells of that cell type further than 30 microns from the nearest leukemia cell (**Figure 5A**).

**Figure 5:**
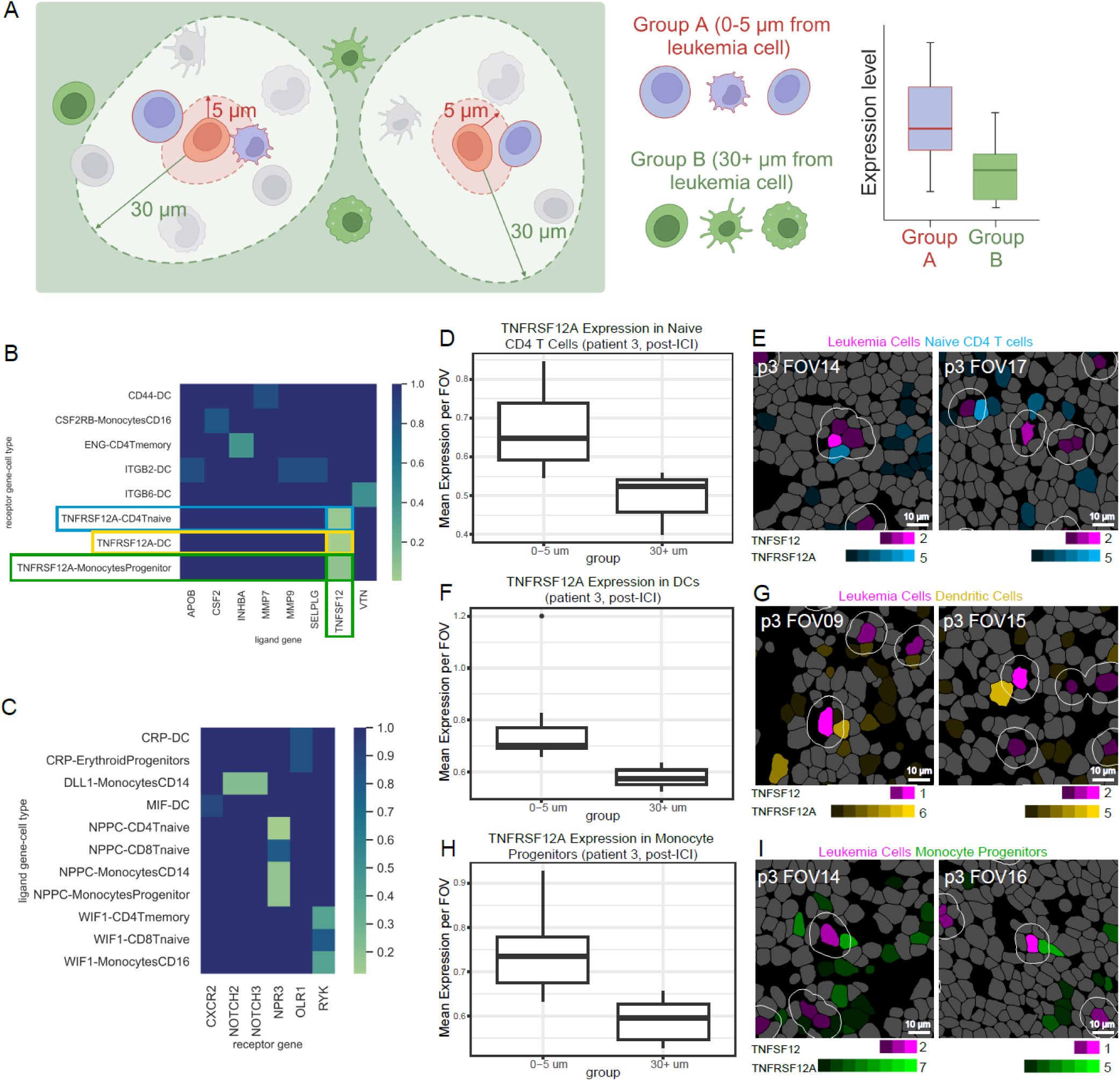
Ligand-receptor analysis. **(A)** Schematic illustrating how the pseudo-bulk method was used to separate cells into “close” (0-5 microns to the nearest leukemia cell) and “far” (30+ microns to the nearest leukemia cell) groups. **(B)** Heatmap of ligand-receptor pairs where the ligand gene is expressed by the leukemia cell (x axis) and the receptor gene is expressed by another cell type (y axis) in patient 3. P-value was calculated based on difference in expression level of the receptor gene between a given cell type’s close and far groups. The two pairs with significantly higher receptor expression in the close group (p-value < 0.05) were TNFRSF12A expressed by DCs/TNFSF12 expressed by leukemia cells and TNFRSF12A expressed by naïve CD4 T cells/TNFSF12 expressed by leukemia cells. The pair TNFRSF12A expressed by monocyte progenitor cells/TNFSF12 expressed by leukemia cells (p=0.07). **(C)** Heatmap of ligand-receptor pairs where the receptor gene is expressed by the leukemia cell (x axis) and the ligand gene is expressed by another cell type (y axis). P-value was calculated based on difference in expression level of the ligand gene between a given cell type’s close and far groups. No significant pairs of biological interest were identified. **(D)** Boxplot comparing the mean expression per FOV of receptor gene TNFRSF12A of all naïve CD4 T cells in the close vs far groups from patient 3 at the post-ICI time point. **(E)** Representative images of segmented cells showing naïve CD4 T cells expressing TNFRSF12A within a 5-micron distance of a leukemia cell expressing TNFSF12. Leukemia cells are in magenta and CD4 T cells are in blue. Reads per cell of each transcript are encoded by dimness/brightness of each color. Scale bars represent 10 microns. **(F)** Boxplot comparing the mean expression per FOV of receptor gene TNFRSF12A of all DCs in the close vs far groups from patient 3 at the post-ICI time point. **(G)** Representative images of segmented cells showing DCs expressing TNFRSF12A within a 5-micron distance of a leukemia cell expressing TNFSF12. Leukemia cells are in magenta and DCs are in yellow. **(H)** Boxplot comparing the mean expression per FOV of receptor gene TNFRSF12A of all monocyte progenitors in the close vs far groups from patient 3 at the post-ICI time point. **(I)** Representative images of segmented cells showing monocyte progenitor cells expressing TNFRSF12A within a 5- micron distance of a leukemia cell expressing TNFSF12. Leukemia cells are in magenta and monocyte progenitor cells are in green.

Ligand-receptor pairs found to be significantly enriched were all observed at the post-ICI treatment time point (**Figure 5B**). Most notably, leukemia cells in patient 3 showed elevated expression of *TNFSF12*, which codes for a proinflammatory cytokine in the tumor necrosis factor (TNF) family called TWEAK, or TNF-like Weak Inducer of Apoptosis (also known as APO3L, CD255). This cytokine can trigger apoptosis in neighboring cells. Surprisingly, the nearby naïve CD4 T cells (**Figure 5D-E**) and DCs (**Figure 5F-G**) had significantly higher expression of *TNFRSF12A*, which codes for the corresponding receptor TWEAKR (also known as FN14, CD266) than that of the same cell types at least 30 microns away from the closest leukemia cell. Monocyte progenitor cells also expressed elevated levels of *TNFRSF12A* (**Figure 5H-I**) with a p-value trending towards significance (p=0.07).

Prior studies have reported that TWEAK binding stimulates activation of the NF-κB (nuclear factor kappa B) pathway in neighboring TWEAKR^+^ cells, leading to TNF-α production and pro-apoptotic autocrine signaling^29,30^. Studies have shown that TWEAK signaling can be particularly lethal to monocytes, promoting monocyte clearance alongside TRAIL and Fas^30^. In a patient with progressive disease, the expression of *TNFSF12* in tumor cells indicates potential leukemia- instigated apoptosis of neighboring myeloid cells. In addition to promoting apoptosis, the TWEAK ligand can also enhance cell proliferation while inhibiting differentiation^31^. This has been observed in myoblasts which were stimulated by TWEAK to proliferate but failed to differentiate^32^. This TWEAK function is particularly interesting in our case, as one of the cell types expressing the receptor TWEAKR was monocyte progenitor cells. It is possible that the TWEAK ligand from leukemia cells could be inhibiting the immune response by dysregulating monocyte progenitor cells from differentiating into functional monocytes.

## Discussion

In this study, we combined the power of traditional scRNA-seq with cutting-edge spatial transcriptomics technology, enabling us to gain insight into changes to the tumor microenvironment of R-AML induced by treatment with an ICI and hypomethylating agent. After immunotherapy, we observed remodeling of the AML bone marrow niche: responders became more enriched with CD14^+^ monocytes, while DCs became more prevalent in nonresponders. Although we did not find a statistically significant enrichment of CD8 T effector cells from baseline to the post-ICI time point, it has been established that in an effective CD8 killing response, CD8 T cells undergo apoptosis after executing the killing function^33^. It is therefore unlikely that CD8 T cells would have been prevalent at the time of the second bone marrow biopsy (8 days after ICI administration). The density shift algorithm identified patient-specific spatial patterns in the cell distribution around leukemia cells before and after ICI treatment. Given that spatial transcriptomics data captures both a cell’s position and its transcriptome, we examined potential ligand-receptor gene pairs being expressed in neighboring cells. Although limited by our sample size, CosMx data showed potential post-ICI TWEAK signaling between leukemia cells and adjacent DC, naïve CD4 T cells, and monocyte progenitor cells.

While spatial transcriptomics technology continues to develop, for the time being, some intrinsic limitations remain. Here we used the CosMx 1000-plex gene panel, which can detect up to 960 different transcripts-still much less than that of traditional scRNA-seq^34,35^. Hence, not all interactions between tumor and immune cells are fully captured due to suboptimal gene coverage. Moreover, spatial transcriptomic technologies capable of detecting transcripts in thicker samples will further enhance detection capabilities. In addition, protein staining of lineage and functional cell markers helps with cell type classification, tumor vs. immune for instance. However, finer-grained classification requires multiple protein markers, which is very challenging with the current available spatial transcriptomic technologies. For cells not identifiable through protein staining, the limited number of transcripts captured per cell in our spatial transcriptomics data made accurate cell typing extremely challenging. We thus chose to use a reference-based method to annotate cells by using overlapping marker gene expression signatures across different cell types from 10x scRNA-seq, which offers a more robust result of cell typing compared with the use of spatial transcriptomics data alone. Other platforms for spatial transcriptomics will have different properties with respect to the area they can analyze, the number of transcripts per cell, the ability to stain for multiple proteins and the ability to perform cell membrane staining.

Another intrinsic limitation of spatial transcriptomics methods is that for true single-cell level data, it requires comprehensive cell segmentation across the entire area of interest. Imprecise delineation of cell boundaries leads to the incorrect attribution of liminal transcripts, which obscures the true cell-specific gene expression patterns and hinders the identification of rare or subtle cellular states, especially at the currently low read depth. Moreover, inaccurate segmentation can confound the computational analysis of intercellular spatial relationships. The utility of single-cell level spatial transcriptomic results is therefore dependent on accurate single- cell level segmentation. To this end, we invested a large amount of effort in optimizing the cell segmentation process, which showed significant improvement over the vendor-supplied segmentation result. Our comprehensive training dataset and optimized segmentation approach can serve as a flexible foundation for developing and benchmarking various new cell segmentation models, ultimately enabling more precise and robust spatial analysis of complex tissues. We have found that it is more effective to train specialist models on membrane protein channel images and combine their segmentation results than to rely on generalist models, at least for the time being. A true cell segmentation foundation model, i.e., a model that can accurately segment cell borders across diverse tissue types in the full range of experimental conditions, will require training on an even larger dataset.

In addition to technological limitations, spatial transcriptomics methods are also prone to substantial sampling bias. In our study, we imaged needle biopsies from the bone marrow, which at the outset provided only a limited view of the tumor microenvironment. Within each biopsy, 5 to 15 FOVs sized 0.985 mm by 0.657 mm were selected for imaging, each covering an area of about 0.65 mm^2^. Between these same-sample replicates, we observed variability in both the transcriptomic and imaging data, providing further evidence that one or more FOVs selected for spatial transcriptomic analysis might not be representative of an entire sample. While capturing the entire tissue area would be ideal, this is usually not logistically and technically feasible. In addition to the increased time and resources necessary to acquire additional FOVs, the process of repeatedly flowing reporters into and out of the flow cell can gradually damage the tissue and cause it to float off the slide. We expect the issue of inter-FOV variability to be a substantial issue in the analysis of spatial data.

As a study designed to assess the clinical feasibility of pembrolizumab-decitabine combination therapy in patients with R-AML, this early phase clinical trial consisted of 10 patients. The criteria that patients needed to have CD34^+^ leukemia for inclusion in the spatial transcriptomics analysis limited the cohort size to 6 patients. Among these 6 patients, treatment outcomes varied substantially: one patient had a complete response, another had a partial/complete response, two finished the regimen with stable disease, and two developed progressive disease. Among these vastly different outcomes, we categorized patients into responder (stable disease/response, n=4) and nonresponder (progressive disease, n=2) groups. In the future, it would be valuable to obtain a direct mutation profile of leukemia cells using spatial transcriptomic data, though this remains challenging with the current spatial technologies. Heterogeneity between patients made it difficult to distinguish treatment-dependent effects from natural patient-to-patient variability. A large-scale clinical study would be necessary to further confirm and validate our findings of the immunotherapy-driven bone marrow niche remodeling.

Despite the limitations of spatial transcriptomics technology and the limited sample size, we show that integrating complex single-cell proteogenomic analysis with spatial-temporal multi-omic analysis in patients with AML is feasible and may lead to a deeper understanding of interactions between leukemia cells and immune cells in the bone marrow niche as evidence by our discovery of the shifts in cell composition in leukemia neighborhood and leukemia cell-mediated TWEAK signaling. Applying our spatial-temporal multi-omic analysis method to a larger clinical cohort should reveal the interactions between tumor and immune cells in the tumor microenvironment and identify potential targets for developing better therapeutics.

## STAR Methods

### Patient Cohort

Bone marrow samples were from the pilot clinical study 17H-0026 (PDAML, NCT02996474, approved by NHLBI Institutional Review Board) with 10 adult R-AML patients to test the feasibility of the combination of pembrolizumab and decitabine. All patients consented to the trial. The treatment and sample collection time were previously reported^4^. The baseline data were collected before any drug administration. The second and third time points corresponded to 8 days after the first dose of pembrolizumab just prior to decitabine initiation (post-ICI treatment, C1D8) and the end of cycle 2 (post-combination therapy, EOC2), respectively. Bone marrow samples were also collected from healthy donors within the age distribution of the patients for scRNA-seq as the reference datasets^5^.

### Single-Cell RNA Sequencing Experiments and Analysis

#### Cell Surface Staining with Oligo-Tagged Antibodies

Cryopreserved BMMCs were thawed into RPMI-1640 supplemented with 10% fetal bovine serum, washed twice with centrifugation at 300x*g* for 5 min at 4 °C in a swing-arm rotor centrifuge, and counted with trypan blue staining on an automated hemocytometer. Between 500,000 and 1 million cells were resuspended in 100 μl of phosphate-buffered saline (PBS) with 1% bovine serum albumin (BSA). Cells were incubated with 5 μl Human TruStain FcX FC receptor blocking solution for 5 minutes at 4 °C, after which 1 μl (0.5 μg) each of the 32 TotalSeq-A oligo-tagged antibodies (Biolegend) were added. Cells were incubated with antibodies for 30 minutes at 4 °C. After incubation, cells were washed twice with 1 mL PBS + 1% BSA, filtered through a 40-μm strainer, and counted with trypan blue staining on an automated hemocytometer. Cells were washed a final time with PBS + 0.04% BSA, and cell concentration was adjusted to between 700 and 1200 cells/μL. Cells were kept on ice until acquisition.

#### Single-Cell RNA Sequencing Library Construction

The 10x Genomics 3’v3 Single Cell Profiling platform was used for scRNA-seq. After staining and cell concentration adjustment, cells were loaded onto a Chromium Chip A with master mix, gel beads, and partitioning oil according to the manufacturer’s instructions. The desired number of captured cells was 4000. Reverse transcription (RT) of mRNA into barcoded first-strand cDNA was performed on the emulsion by incubating at 53 °C for 45 minutes in an Eppendorf Mastercycler X50a. After RT, emulsions were broken, and cDNA purified using Dynabeads MyOne SILANE. At this stage, cDNA was amplified by PCR (98 °C for 45 seconds; 13 cycles of 98 °C for 20 seconds, 67 °C for 30 seconds, 72 °C for 1 minute; 72 °C for 1 minute). Additional primers were added to ensure adequate amplification of oligo tags to later generate ADT sequencing libraries. Amplified cDNA was mixed with 0.6X SPRIselect magnetic beads (Beckman Coulter); after magnetic separation, supernatant was transferred for ADT library generation, and the remaining pellet was used for constructing the gene expression (GEX) library. The concentration of amplified cDNA was measured at a 1:10 dilution with a High Sensitivity (HS)-D5000 chip on an Agilent Tapestation.

For the GEX library, 50 ng of amplified cDNA was carried forward for library construction. Fragmentation, end-repair, and A-tailing was performed (32 °C for 5 minutes; 65 °C for 30 minutes), followed by double-sided size selection using SPRIselect beads. Adaptor ligation (20 °C for 15 minutes), cleanup with SPRIselect beads, and PCR amplification with sample indexing primers (98 °C for 45 seconds, 14 cycles of 90 °C for 20 seconds, 54 °C for 30 seconds, 72 °C for 20 seconds; 72 °C for 1 minute) were performed in that order. A final double size selection was again done using SPRIselect beads.

To make ADT libraries, sample index PCR using unique indices was performed with the transferred supernatant taken immediately after initial cDNA amplification (98 °C for 45 seconds; 9 cycles of 98 °C for 20 seconds, 54 °C for 30 seconds, 72 °C for 20 seconds; 72 °C for 1 minute) using Kapa HiFi Hotstart Readymix (Kapa), followed by a single-sided size selection cleanup with SPRIselect beads.

#### Single-Cell RNA Sequencing Library Quality Control and Sequencing

GEX and ADT libraries were run on an HS-D5000 chip using an Agilent Tapestation at a 1:10 dilution, and the average size of all libraries was calculated (GEX libraries were around 500 base pairs [bp] in size; ADT libraries were about 200 bp). The libraries were quantitated in triplicate using qPCR (Kapa) at 4 dilutions from 1:40,000 to 1:5,000,000 to calculate nanomolar (nM) library concentrations. The scRNA-seq libraries were then pooled in the appropriate ratio so that each library would be sequenced at the correct depth and sequenced on an Illumina HiSeq 3000 using read lengths of 150 bp from read 1, 8 bp of the i7 index, and 150 bp from read 2. Target sequencing depth for the GEX libraries was 50,000 read pairs per cell and 5,000 read pairs per cell for ADT libraries.

#### Single-Cell Data Preprocessing, Integration, and Clustering

Sequencing libraries were preprocessed using Cell Ranger version 3.1.0 (10x Genomics) to output filtered UMI matrices for downstream analysis^36^. The number of barcode mismatches was set to be 0 to minimize demultiplexing error. Cells with overlapping cell barcodes between GEX and ADT results were included in the multimodal analysis.

Seurat version 3 was used for sample basic quality control, normalization, and integration^37^. Data were normalized based on regularized negative binomial regression and 2000 variable features were selected for each sample based on a variance stabilizing transformation. Anchor cells, which are pairs of cells in matched biological states, were identified across samples to integrate and match the overlapping cell populations. Integrated datasets were scaled, and the number of principal components was chosen to represent the heterogeneity in gene expression. The cell clusters were constructed by applying a graph-based clustering approach, and the data were explored by UMAP for visualization. Highly expressed proteins were identified for cell type annotation. Expression of several categories of genes including exhaustion, cytotoxicity, and naïve signatures were extracted to calculate the respective gene scores for each cell type. Leukemia-associated clusters were labeled based on markers from previous clinical flow cytometry records and the labeling of cell clusters was confirmed by performing clustering analysis on proteins for each patient sample. Protein-only clustering done separately for each sample was also used to confirm other major cell types such as T cells. The InferCNV^38^ pipeline was implemented to identify cells with a known copy number variation for samples from patients with reported cytogenetic abnormalities. Malignant cell populations identified using different methods were compared with each other by calculating the overlap percentages and using the

joint UMAP for visualization.

### Spatial Imaging Processing with Proteomic and Transcriptomic Profiling

#### Sample Preparation

Formalin-fixed paraffin-embedded slides prepared from bone marrow biopsies at time points A (pretreatment), B (post pembrolizumab), and C (post both pembrolizumab and decitabine) from six patients were analyzed using a pre-commercial CosMx SMI (NanoString Technologies Inc). All three biopsy samples from a given patient were placed on the same glass slide (**Figure 1A**). To improve tissue adherence, the slides were baked overnight (60 °C). Next, the Leica Bond RX system was used to expose RNA targets on the samples through a process of deparaffinization, proteinase K digestion (3 µg/mL, ThermoFisher; incubated at 40 °C for 30 minutes), and heat- induced epitope retrieval (using Leica buffer ER1 at 100 °C for 15 minutes). Samples were then rinsed twice with diethyl pyrocarbonate (DEPC)-treated water and incubated for 5 minutes at room temperature with fiducials (Bangs Laboratory) diluted 1:1000 in a 2X SSCT solution (2X saline sodium citrate with 0.001% Tween 20). After excess fiducials were removed by rinsing the samples with 1X PBS, the samples were fixed for 5 minutes at room temperature in 10% neutral buffered formalin, then rinsed for 5 minutes each with Tris-glycine buffer (0.1 M glycine, 0.1 M Tris base in DEPC-treated water) and 1X PBS. Next, samples were blocked for 15 minutes at room temperature with 100 mM *N*-succinimidyl acetate (NHS-acetate; ThermoFisher) in NHS- acetate buffer (0.1 M NaP, 0.1% Tween at pH 8 in DEPC-treated water). After a 5-minute 2X SSC rinse, samples were protected with an Adhesive SecureSeal Hybridization Chamber (Grace Bio-Labs) before overnight hybridization with RNA in situ hybridization (ISH) probes. The RNA ISH probe panels included 960 probes and 19 negative control probes. To prepare the ISH probe mixture, RNA ISH probes were denatured for 2 minutes at 95 °C, then placed immediately on ice. The probe mix was then pipetted into the hybridization chamber enclosing the sample and covered with adhesive tape to prevent the hybridization solution from evaporating during the 37 °C overnight hybridization. After hybridization, samples were washed twice with a 50% formamide (VWR) solution in 2X SSC, each for 25 minutes at 37 °C. They were next rinsed twice for 2 minutes in 2X SSC alone at room temperature. Before the samples were encased on each slide in a custom flow cell, they were blocked for 15 minutes with 100 mM NHS-acetate and washed twice for 2 minutes in 2X SSC alone at room temperature.

#### Data Acquisition

The SMI RNA detection and imaging procedure is described by He et al^15^ and summarized here. The prepared flow cell was loaded into the SMI, where several rounds of reporters would be cycled in, imaged, and removed to detect the presence of all transcripts in the panel. Before beginning the cycling, reporter wash buffer (1X SSPE, 0.5% Tween 20, 0.1 U/µL SUPERase•In RNase Inhibitor at 20 U/µL, 0.1% Proclin 950 and DEPC-treated water) was flowed into the flow cell to rinse the samples and remove any air bubbles. Next, the SMI performs a preliminary scan of the whole area covered by the flow cell, at which point 5 to 10 FOVs were selected from each biopsy that would be captured by the camera in the coming reporter cycles. To begin the first RNA readout cycle, 100 µL of reporter pool 1 flowed into the flow cell and was left to incubate for 15 minutes before 1 mL of reporter wash buffer flowed in to remove any unbound probes. Before the instrument imaged the fluorescent probes, the wash buffer was replaced with an imaging buffer (80 mM glucose, 0.6 U/mL pyranose oxidase from *Coriolus* sp., 18 U/mL catalase from bovine liver, 1:1000 Proclin 950, 500 mM Tris-HCl buffer pH7.5, 150 mM sodium chloride, and 0.1% Tween 20 in DEPC-treated water). The SMI captured 9 z-stacked images, at a step size of 0.08 µm, from each FOV. After imaging, the fluorophores on the reporter probes were UV cleaved and washed away with strip wash buffer (0.0033×SSPE, 0.5% Tween 20 and 1:1000 Proclin 950). This reporter pool, imaging buffer, and wash buffer cycle repeated 15 more times for a total of 16 cycles total (once per reporter pool). In total, 9 rounds of this 16-cycle process were repeated. The last step was to capture sample morphology. The samples were incubated for 1 hour with a solution of fluorescent-labeled antibodies, which included: CD298 at 1:40 dilution (Abcam, EP1845Y), B2M at 1:40 dilution (Abcam, EP2978Y), CD34 at 1:40 dilution (Abcam, Qbend/10; conjugated to AF647), and DAPI. One additional protein marker, CD3 at 1:20 dilution (Abcam, F7.2.38), was not directly conjugated to a fluorophore, so the 1 hour of CD3 incubation was followed by 1 hour of donkey anti-mouse AF594 incubation (1:80 dilution). After excess antibody was washed away with reporter wash buffer and the wash buffer was replaced with imaging buffer, the instrument captured 9 z-stacks per FOV.

#### Image Preparation: Normalization & Deconvolution

Of the 5 channels imaged, only 2 were needed for cell segmentation: DAPI for nucleus segmentation and CD298/B2M for membrane segmentation. To create a crisper image for image annotation, we deconvolved the DAPI and CD298/B2M channels using Huygens Professional version 22.04 (Scientific Volume Imaging, The Netherlands, http://svi.nl). Finding the cell boundaries easier to detect but the nuclei dimmed, we layered the deconvolved DAPI channel with the raw DAPI channel. Next, both the deconvolved CD298/B2M channel and hybrid raw/deconvolved nucleus channel were clipped to a range between the 5th and 99.99th percentiles of their raw grayscale values before each protein channel was mapped to its RGB color channel (DAPI in blue, CD298/B2M in green).

#### Model Training

We used the Cellpose^18^ train-your-own model feature to train 2 models: one for nucleus segmentation and one for cell membrane segmentation (using the B2M/CD298 membrane marker). Using manual annotation, we first generated a total of 448,439 ground truth nucleus masks. Once enough nuclei had been annotated to train our own model, we switched to a semiautomatic annotation approach. We used our trained nucleus model to predict nucleus masks and manually corrected them. We used the same approach to generate a membrane prediction model, with 318,401 ground truth membrane masks used in the final dataset.

#### Implementing Segmentation Models

Of the 142 FOVs imaged, nuclei masks were manually corrected on 76 and membrane masks on 22. For the remaining FOVs, our trained nucleus and membrane models were used to generate predicted masks of each respective class. Wanting to have a single mask to represent each cell in (including both nucleus and membrane), we merged nucleus and membrane masks that corresponded to the same cell for all FOVs. When evaluating the output of the trained model in comparison with the ground truth, we computed an F1 score using the formula:

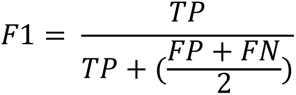

Where TP is the number of true positives, FP is the number of false positives, and FN is the number of false negatives. To compute this metric, we used an IOU threshold of 0.7, meaning that for a single cell label to be considered a true positive, it must overlap with a ground truth label by 70% or more. Any less overlap and that label would count as a false positive.

#### Cell Type Identification for Spatial Transcriptomics Data

Before proceeding with assigning cell types by gene expression profile, we identified leukemia cells using an approach based on cell morphology and CD34 protein expression. Raw CD34 channel intensities were clipped below the 5th percentile and above the 99.9th and then normalized from 0 to 255 for each FOV. Individual images of every cell’s normalized CD34 protein channel were resized to 224x224 pixels and triplicated to create a pseudo-RGB image. For each patient, the pseudo-RGB images of all cells imaged from that patient were fed into the EfficientNet classifier^20^. The vector of 672 features extracted from each image in the B0 layer was then used to cluster the cells via UMAP. Overlaying the mean CD34 protein intensity expressed in each cell on the UMAP revealed, as anticipated, an intensity-dependent clustering. Since cell shape is also encoded in the features of each image, leukemia cells clustered separately from other CD34^+^ cells with different morphology (chiefly endothelial cells of the blood vessels).

RBCs, which are morphologically distinguishable from other cells owing to their lack of nucleus and distinct shape, were identified and removed before transcript-based cell type assignment. Like the process used to identify leukemia cells (but using normalized RGB images that contained the DAPI, B2M/CD298, and CD34 channels), single-cell image features extracted from the B0 layer of the EfficientNet^20^ classifier were used to cluster cells by morphology. Representative cells from each cluster on the UMAP were visualized to identify which clusters contained RBCs. All cells of suspected RBC clusters were highlighted on each FOV and reviewed to verify that the RBCs had indeed been identified. This process was completed independently for each patient.

Total transcript count was used as the quality control metric to remove low-quality cells (<20 transcripts assigned to the identification), and 3 FOVs with less than 300 cells remaining were removed from the analyses. InSituType^38^, a likelihood-based cell typing method, was used to annotate the remaining cells. Cell types identified in 10x single-cell experiments were included in the supervised cell typing pipeline, and the marker genes were selected based on prior knowledge and a differential expression list to create the reference profiles. Because the data from different samples were heterogeneous, the following cell type annotation steps were performed within each sample so that the sample variability would not impact the annotation result. The expression matrices for all cell typing marker genes were extracted and the average expression of each cell type was calculated for each cell. Reference cells were selected by choosing those with only one cell type marker expressed, and the reference profiles were created by taking the average expression of all genes from cells labeled as reference cells for each cell type. The reference matrix was the input for the algorithm with genes in rows and cell types in columns. Average negative control values were provided to adjust the count and provide the normalized count, and the posterior probability was used to determine the cell types. The first round of the analysis determined the broad cell types, and an additional round of creating reference and calculating posterior probability was performed for immune cell subtypes to obtain more specific annotation for B cells and T cells. The average expressions of marker genes for the final cell types were compared using a heat map to confirm the accuracy of the gene expression profile of each population. External validation was performed by using the cell size information obtained from the cell segmentation step.

### Spatial Distance Measurement and Cell Distribution Modeling

#### Distance Determination

The spatial genomic data contained local coordinates of all transcripts within each FOV. Each cell, however, was represented as a polygon, and cell-to-cell distances were found by computing the minimum distance between the edges of every pair of cells in each FOV.

#### Generalized Linear Mixed Effect Model for Spatial Distribution

For each leukemia cell, every other cell was labeled with a distance group by how far it was to this leukemia cell, with 0 representing directly touching and every other group labeled by increasing distances of 5 microns. Then we summarized the number of cells for each cell type around this leukemia cell. The input for the Poisson generalized linear mixed model was the count for different cell types across distance groups for each leukemia cell.

Because we were interested in testing whether there were significant changes in any cell types across patient response and time points, one Poisson generalized linear mixed model^39^ was fit for each cell type, with the count for the cell type of interest as the response variable; the 3-way interaction of patient response (responder/nonresponder), sample collection time points, and distance group as the main effects; and FOVs as a random effect. Because the total cell count of each FOV across different distance groups strongly correlated with the count of each cell type in that region, we considered this total cell count as an offset of the model.

P-values were recalculated using the fitted model for different combinations of coefficients of interest to investigate the difference for responders between time points, and the difference between responders and nonresponders at baseline. Bonferroni correction was used to generate the adjusted p-values considering all models fitted to determine any significant changes, with two-sided adjusted p-values of 0.05 as statistically significant.

#### Cell Type Density Shift

To examine the spatial distribution shift between time points, we started with defining the minimum distance of one cell to another cell type within one FOV as the shortest distance between the cell and any cells of the other cell type. For example, if the two cell types of interest were leukemia cells and T cells, suppose for one FOV, there were 𝑀 leukemia cells 𝑚_1_, 𝑚_2_, …, 𝑚_𝑀_; 𝑁 T cells 𝑛_1_, 𝑛_2_, …, 𝑛_𝑁_. The pairwise distance matrix for all leukemia cells and T cells was defined as:

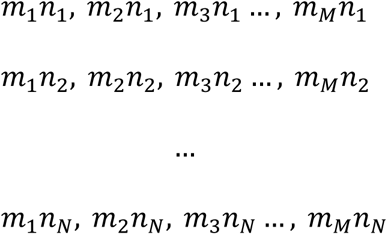

The minimum distance of one T cell 𝑛_𝑖_ to leukemia cells was defined as the minimum value of that row of distance values:

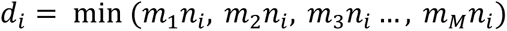

For all the T cell 𝑁 values in this FOV, the distribution of minimum distance was estimated by getting the minimum value of each row in that matrix:

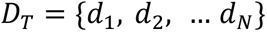

The distribution of 𝐷_𝑇_was the minimum distance distribution from T cells to leukemia cells for this FOV. We considered the FOVs from one time point of one patient as sample replicates, and the distribution of one sample was the aggregated results from all FOVs:

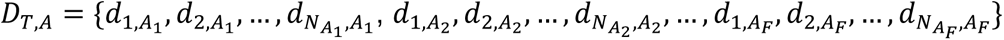

For one patient sample at time point 𝐴, with 𝐹 numbers of FOVs from this sample, and for FOV 𝑓 ∈ {1, 2, …, 𝐹}, there were 𝑁_𝐴𝑓_ number of T cells.

To create the background distribution for the density shift test, we started with constructing the background distribution of any cells to leukemia cells. Similar as what was described, for the same FOV with 𝑀 leukemia cells, 𝐾 other cells including 𝑁 T cells (𝑁 < 𝐾), the background distribution from a random cell group of the same number of cells to leukemia cells was obtained by 100 random permutations. For each permutation, we randomly sampled 𝑁 cells from 𝐾 cells: 𝑛′_1_, 𝑛′_2_, …, 𝑛′_𝑁_ and obtained the following pairwise distance matrix:

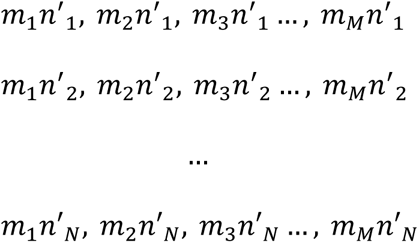

The same definition was used to get the background distribution 𝐷^′^ for one sample.

For 2 time points of interest 𝐴 and 𝐵 (here we considered baseline as time point 𝐴, and the post- ICI time point as time point 𝐵), we obtained the distributions 𝐷_𝑇,𝐴_ and 𝐷_𝑇,𝐵_, and used the 𝑑𝑒𝑛𝑠𝑖𝑡𝑦 function in R to get the respective probability density functions 𝑝𝑑𝑓_𝑇,𝐴_ and 𝑝𝑑𝑓_𝑇,𝐵_ for 𝐷_𝑇,𝐴_ and 𝐷_𝑇,𝐵_, with the same distance range 𝑥 = (𝑥_1_, 𝑥_2_, …, 𝑥_𝑙_, …, 𝑥_𝐿_).

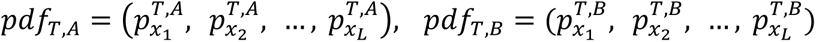

The density shift distance from time 𝐴 to time 𝐵 was defined as

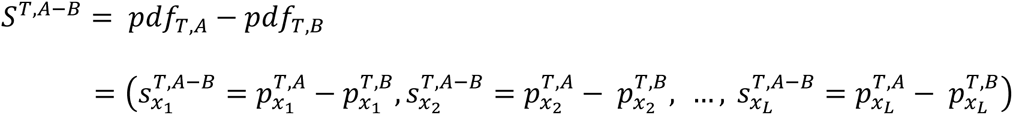

With the same definition, for any random permutation 𝑗𝑗 (𝑗𝑗 = 1, 2, …, 𝐽), we could obtain its density shift distance:

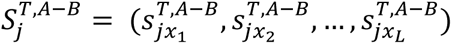

To determine whether the density shift was significant, we checked for each 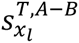, where it fell with regard to the distribution of 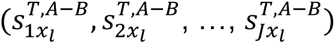. The shift was determined to be significant if the data 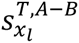 were outside of 1 standard deviation of the simulated background (either direction). For visualization, the color of the shift in density was determined by the magnitude of the shift.

A simulation study was implemented by ordering the cell-to-cell distance and setting the closet or the furthest cell as a T cell to test whether the algorithm could identify the pattern shift. This process was repeated for all cell type pairs of interest to identify potential patterns in spatial shift across treatment time.

#### Ligand-Receptor Analysis

The human ligand-receptor gene pairs were downloaded from CellTalkDB^40^ with 3,399 pairs in total and 719 pairs overlapping with the spatial gene panel. For each cell other than leukemia cells, the distance between the cell and its nearest leukemia cell was calculated. Two cell groups were extracted for the analysis, one with minimum distance less than or equal to 5 microns (close group), and the other with minimum distance greater than or equal to 30 microns (far group). Because the data for each FOV were sparse, we adopted a bulk approach to study potentially different ligand-receptor pairs. The expression data were aggregated for each FOV of each sample (patient and time point) by calculating the mean expression of genes in the ligand-receptor reference for each cell type. We analyzed the data 2 ways by considering genes from the ligand list on the leukemia cells or from the receptor list on the leukemia cells. For each way of considering the gene list, the median expression level of the genes was aggregated for each cell type within one FOV and genes highly expressed in leukemia cells were defined as those with median expression greater than 2 standard deviations away from the expression of other cell types. To test the other gene list on other cell types, Mann-Whitney U test was used to compare the median expression level of genes between 2 distance groups (far group vs close group) within each sample for different cell types respectively considering FOVs as replicates. Ligand receptor results were summarized based on the 2 different tests on ligand and receptor gene lists, and adjusted p-values are reported.

## Software

R version 4.3.2 was used for the analyses and figure creation. Python 3 was used for spatial transcriptomic analyses (minor version varied by virtual environment). Illustrations were created with BioRender.com and Adobe Illustrator.

## Data and Code Availability

Patient clinical data were published in the previous paper^4^. Sequencing data are available for download at GEO (GSE271406) and the CosMx images with annotations on Zenodo at https://zenodo.org/doi/10.5281/zenodo.12730053 upon acceptance. The code for analyzing the data is at the GitHub repository https://github.com/chenzhaolab2023/AML-Spatial.

## Acknowledgment

The study is supported by the Intramural Research Programs of the NCI Center for Cancer Research, NIH Clinical Center, and NHLBI, respectively. This work was supported in part by the Division of Intramural Research, NIAID, NIH. The study was also supported in part by the National Institute of General Medical Sciences of the National Institutes of Health under award number R35GM149323. This work utilized the computational resources of the NIH HPC Biowulf cluster. We thank Drs. Ronald N. Germain and Lichun Ma for valuable advice.

We thank the NanoString customer experience and bioinformatics teams for technical support. Parts of the figures are used with the permission of NanoString Technologies, Inc.

## Author contributions

Conceptualization, G.G., C.H., and C.Z; methodology, G.G., M.B., K.H., and C.Z.; data curation, G.G., M.B., A.W., E.M., J. R., S. K., A. B., K.H., C.Z.; formal analysis, G.G., M.B., H. D., G. Z., E.S. ; funding acquisition, C.H., K.H., C.Z.; investigation, G.G., M.B., K.H., C.Z.; project administration, P.D., C.H., K.H., C.Z., J.H.; resources, C.H., K.H., C.Z.; supervision, K.H., C.Z.; validation, G.G., M.B., K.H., C.Z.; visualization, G.G., M.B., M.G.; writing - original draft, G.G., M.B.; writing - editing, J.H., K.H., C.Z.; writing - reviewing, L.D., M.G.,.

## Declaration of Interests

J.D., S.K., A.B., and P.D. are employees and stockholders at NanoString Technologies Inc. All other authors declare no competing interests.

## Supplemental Information

Supplement S1 [Figures S1-S11] and Tables S1-S3.

## Notes

https://github.com/chenzhaolab2023/AML-Spatial

