## Supplemental figures for "Single Cell Spatial Transcriptomics Reveals Immunotherapy-Driven Bone Marrow Niche Remodeling in AML"

### Supplementary Information: Figures S1-S9

**Figure S1. Top variable genes and proteins across study time points.** No significant patterns were identified by comparing different time points or patient responses. **(A)** Top variable genes across all bone marrow samples from patients. **(B)** Top variable protein targets across all bone marrow samples from patients. **(C)** Top variable protein targets across all blood samples from patients.

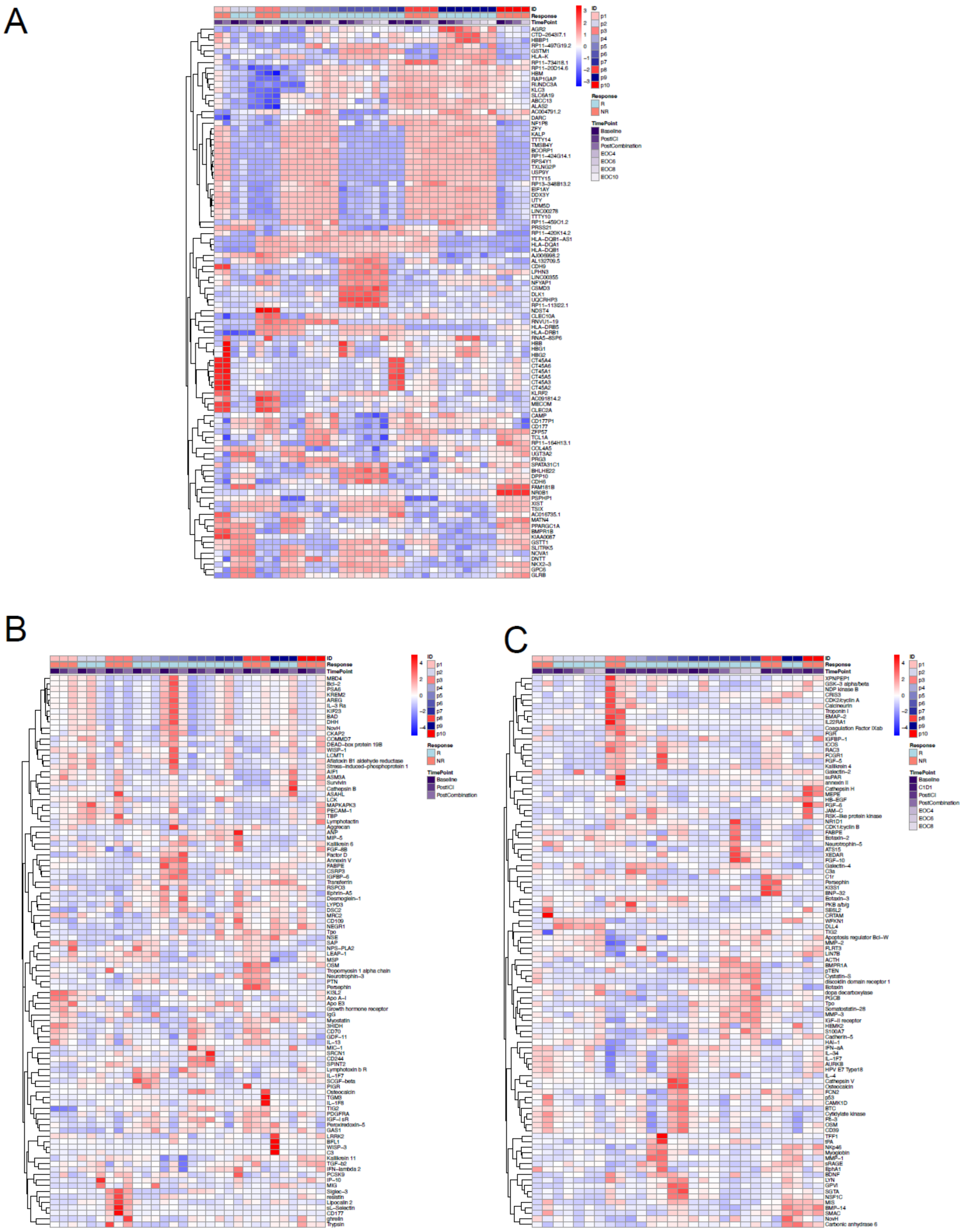

**Figure S2. Cell type distribution and annotation details for 10x single-cell RNA sequencing. (A)** Cell type distribution across all samples. **(B)** UMAP visualization for all samples and by group for patients or healthy donor (HD). **(C)** Leukemia cells identified with CD34/CD117 protein expression overlapped with patient samples. **(D)** Most mature B cells were from healthy donor samples with high CD19/CD22 protein expression. **(E)** Patient-specific leukemia cell clusters. **(F)** Copy number variation was identified for patient 3 (left) and patient 6 (right) by InferCNV. Related to Figures 1 and 2.

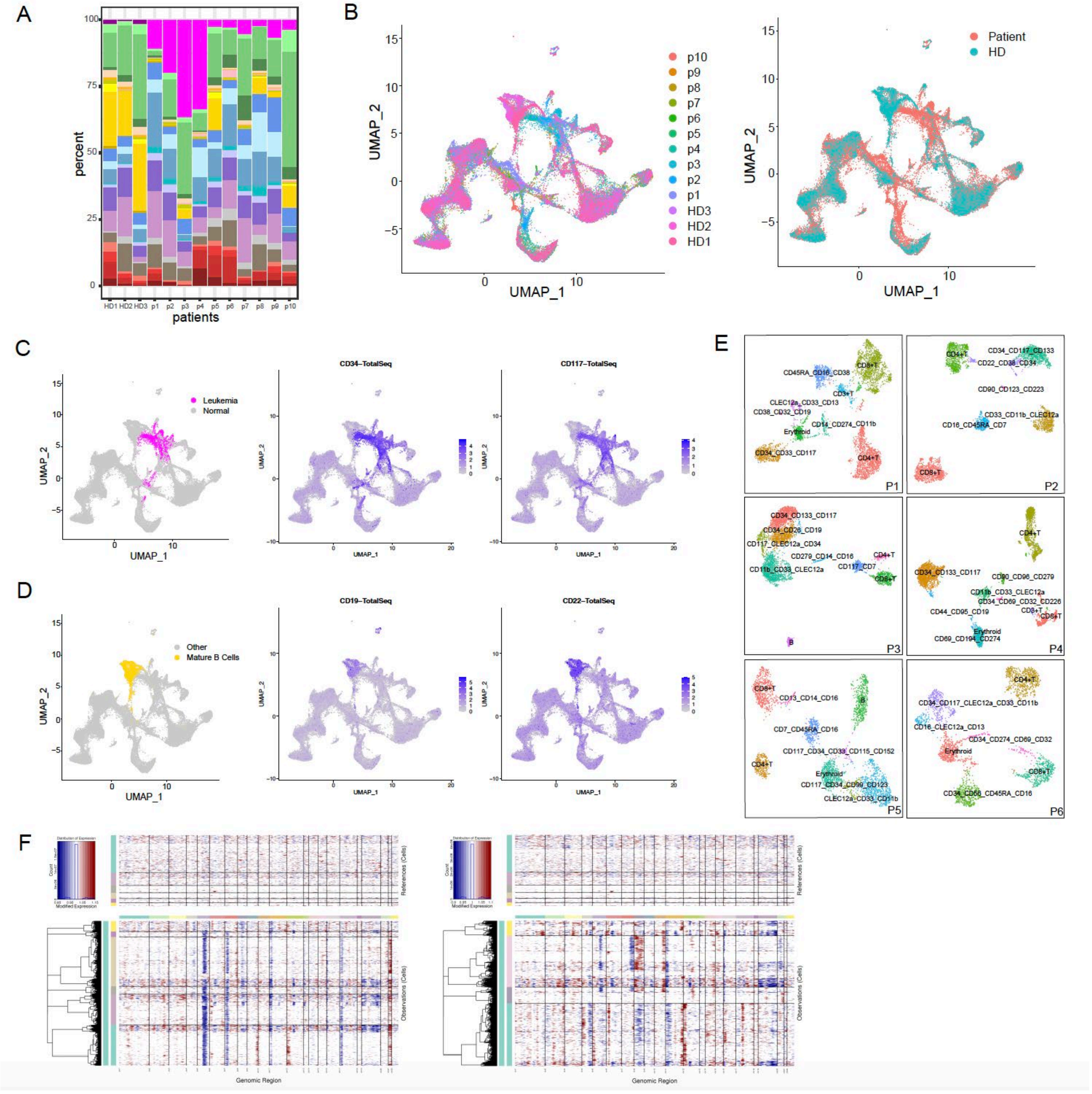

**Figure S3. Training and implementation of cell segmentation models. (A)** A region of one FOV showing two channels: DAPI (blue) and B2M/CD298 (green). We used the **(B)** DAPI and **(C)** B2M/CD298 channels to train nuclear and membrane segmentation models, respectively. For each channel, images were broken up between a training set (90%) and test set (10%). Separately for the **(D)** nuclear and **(E)** membrane channels, we segmented the test set of images using both our trained model and the corresponding pre-trained Cellpose model and evaluated F1 scores across a range of IOU thresholds. **(F)** Schematic to illustrate the process of generating ground truth labels and training each segmentation model. In phase 1, we manually annotated all individual cells in a selected region and used this as ground truth to train untrained Cellpose models. In phase 2, once we had enough manually annotated ground truth images to train a functional model, we used the trained model to predict the cell boundaries on new, unannotated regions. Since the model was still suboptimal, we manually corrected its segmentation and used this as additional ground truth, accelerating the annotation process. **(G)** An illustration of how we merged nuclear and membrane segmentation results to obtain one single label for each cell. Only nuclei without a corresponding membrane were expanded to prevent assigning transcripts to a cell when they were not actually enclosed in that cell. **(H)** We compared the default segmentation provided by NanoString (based on Cellpose) with the fusion of our trained nuclear and membrane segmentation outputs, using our manually annotated data as ground truth. The dark areas in A-C are empty fat droplets in bone marrow.

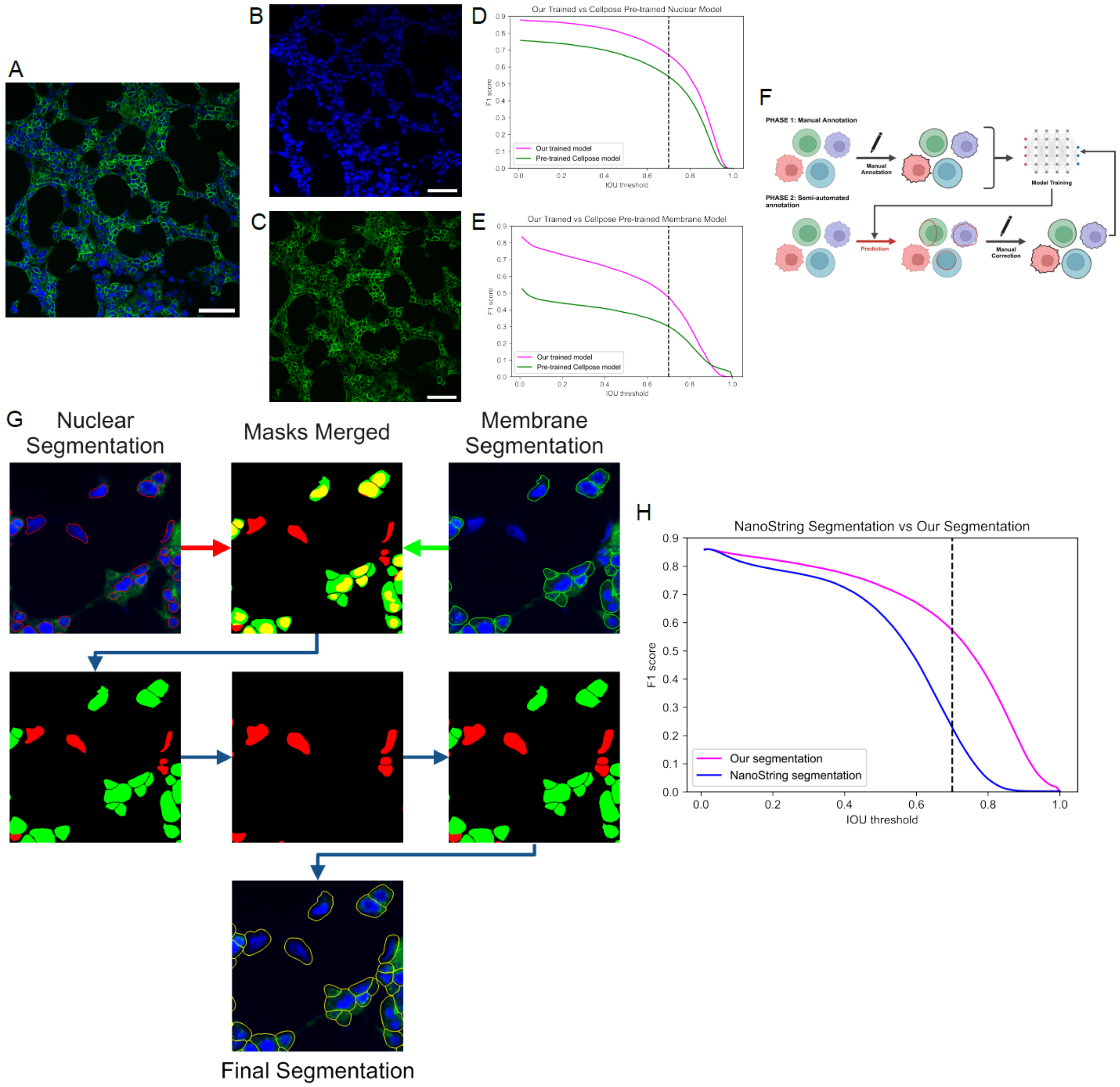

**Figure S4. Cell segmentation generated from CosMx as compared to our custom trained model. (A)** Single cell segmentation result generated from CosMx workflow overlaid in white on a portion of one field of view. DAPI is shown in blue and the PANCK/CD298 membrane marker in green. Scale bar represents 50 µm. **(B)** Segmentation result using our trained model on the same region. A few regions **(1-4)** are highlighted to show the improved performance of our model on this dataset.

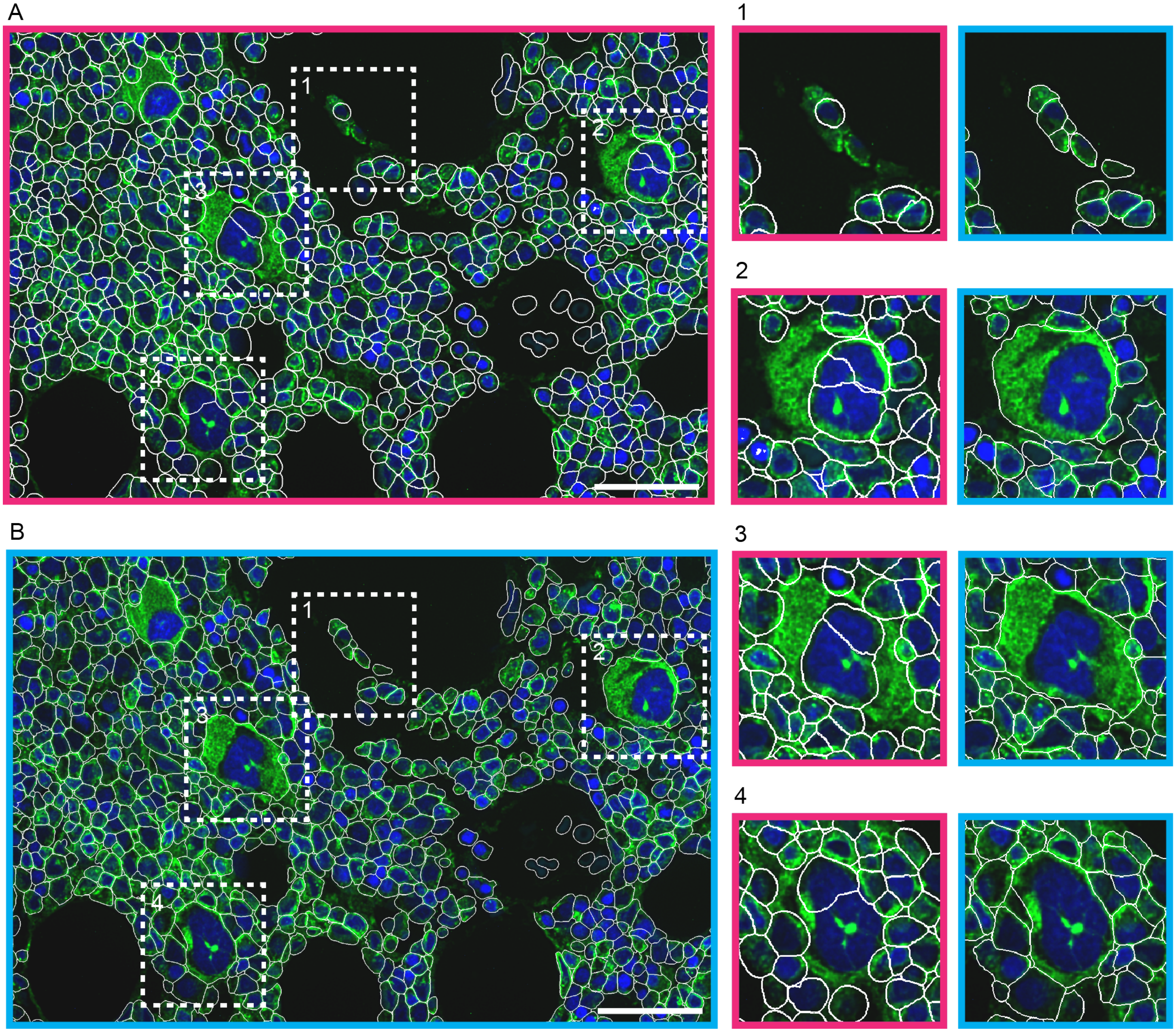

**Figure S5. Quality checking of CosMx data. (A)** Transcript distribution in one example FOV from each patient where each transcript in the panel is given a unique color. Scale bars represent 50 microns. **(B)** Summary of transcript count per cell across all FOVs for each patient. Different colors represent data collection time points, baseline (A, red), post ICI treatment, post combination therapy at end of cycle 2 (C, blue).

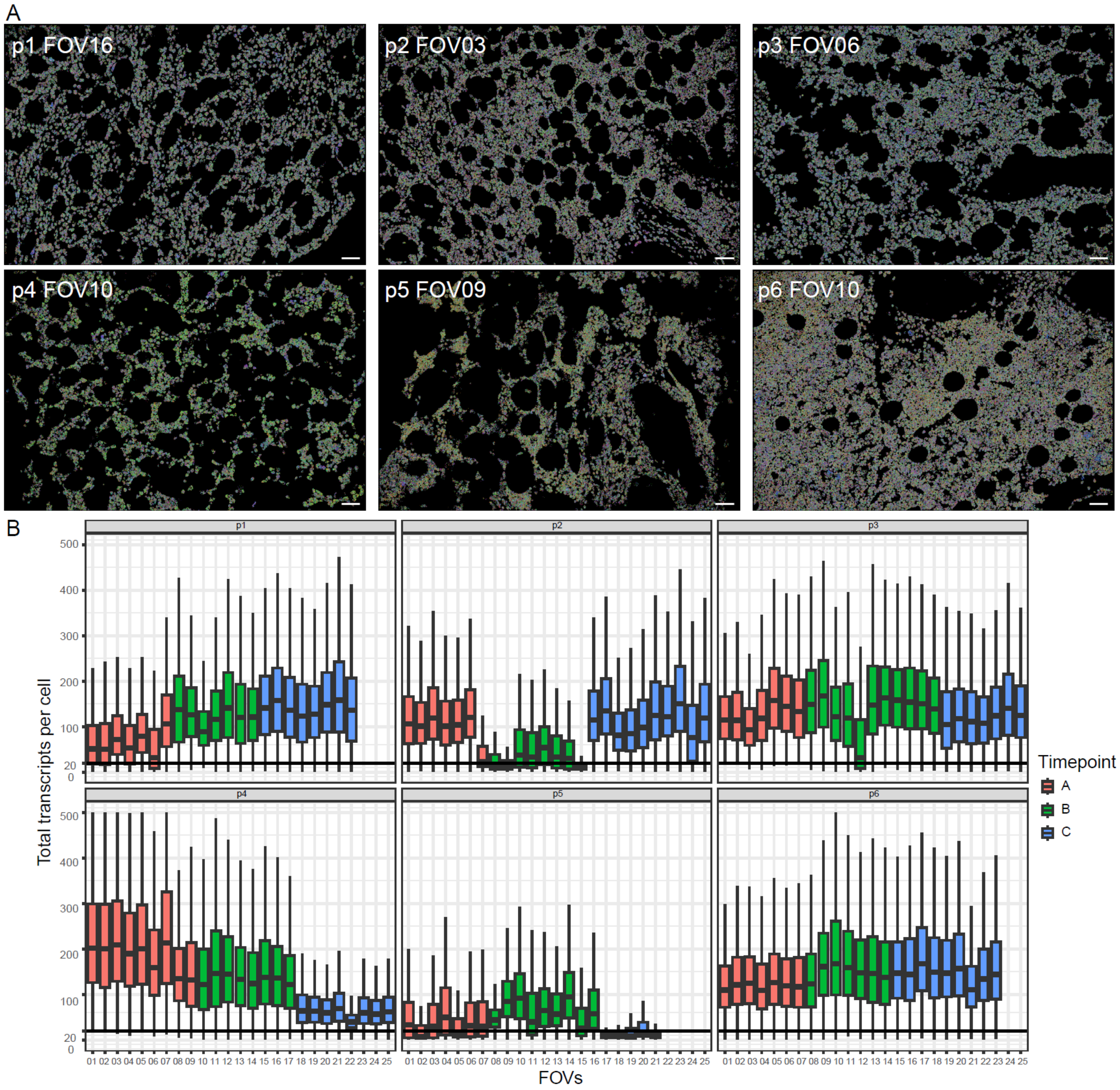

**Figure S6. Identifying red blood cells by morphology for removal. (A)** Portion of one field of view showing DAPI in blue and the PANCK/CD298 membrane marker in green. Scale bars represent 50 µm. **(B)** Single cell segmentation result overlaid on the same image, with red blood cells identified by morphology (via feature extraction and clustering methods) outlined in red.

A

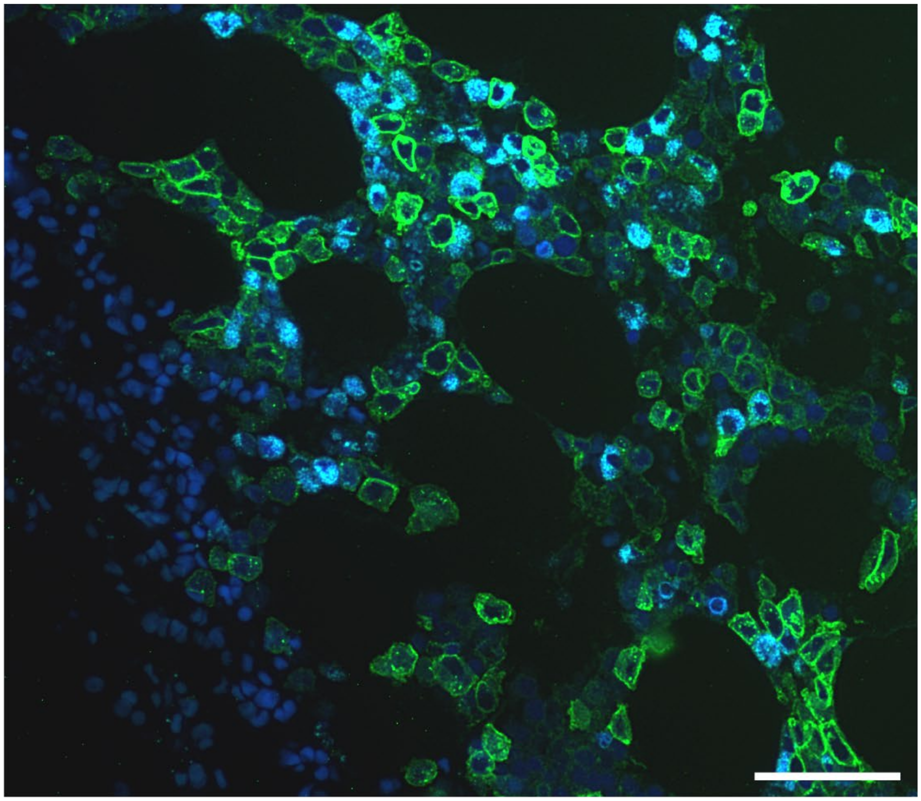

B

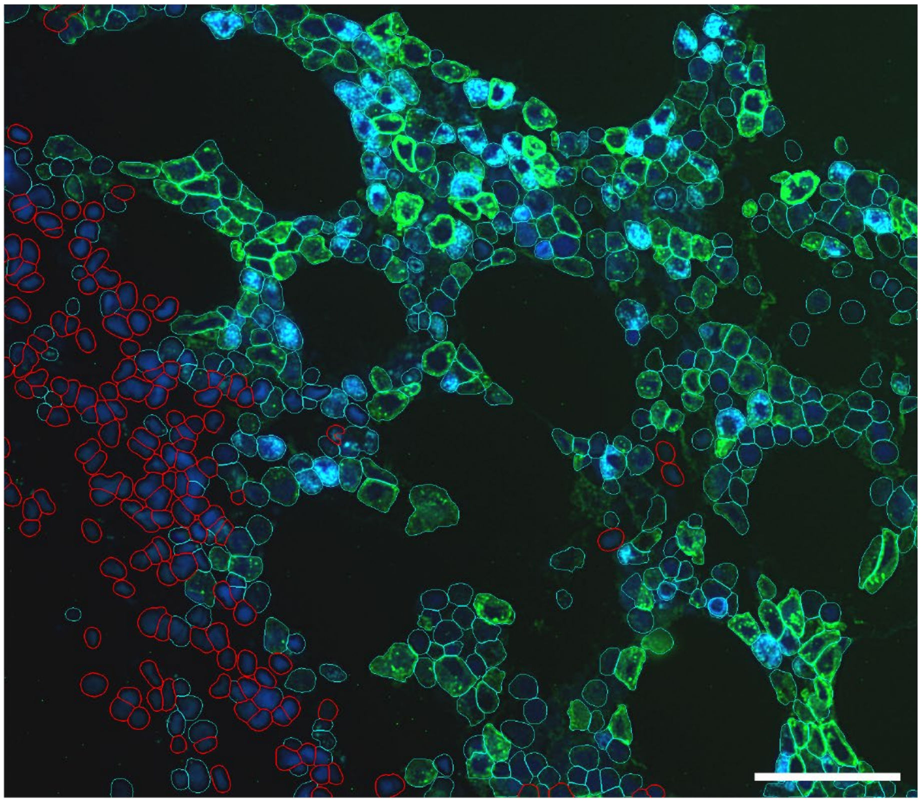

**Figure S7. Leukemia cell calling using morphology and CD34 protein expression. (A)** After segmenting all cells, we created **(B)** single-cell crops of every cell from every FOV, overlayed on a black background of a set size. **(C)** We extracted the CD34 protein channel from each single cell crop, **(D)** yielding a total of 558,515 single-cell CD34 channel cropped images. **(E)** All images were input into the EfficientNet Classifier, saving the B0 layer as a final output. As a preprocessing step, all single-channel, grayscale images were triplicated to create an RGB image and resized to 224x224 pixels (the required format and size for EfficientNet). **(F)** The saved B0 layer was a vector of 672 features for each single-cell image. **(G)** For each patient, we clustered these cells by their CD34 channel features using Phenograph and overlaid the resulting UMAPs with **(H)** CD34 protein expression per cell to identify **(I)** the likely leukemia cell clusters. **(J)** Candidate leukemia cells were viewed in the context of their respective FOVs to verify their identity. All scale bars represent 10 microns.

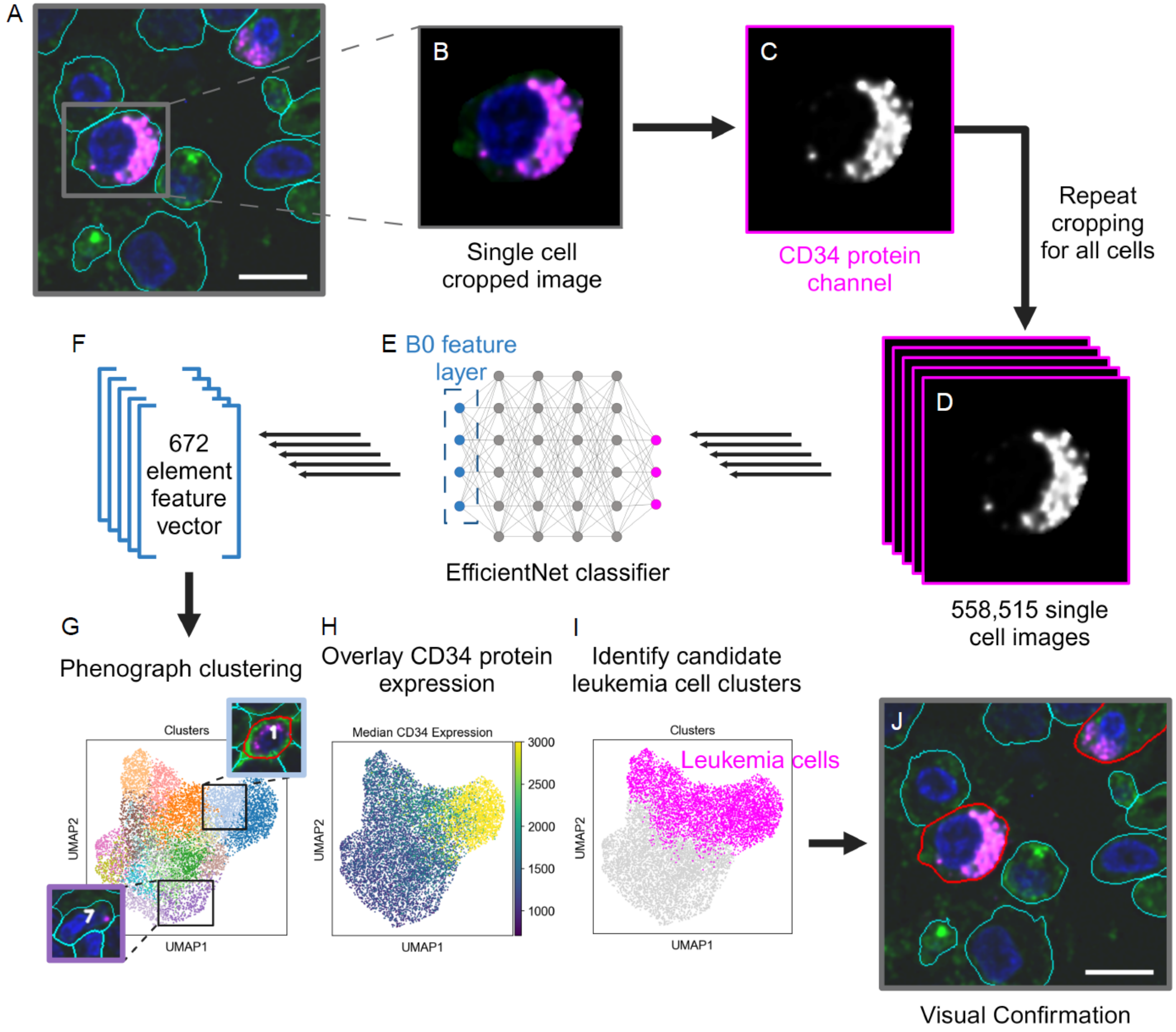

**Figure S8. Cell edge-to-edge distance measurement. (A)** Illustration of measuring cell-to-cell distances using cell edges as opposed to cell centroids. In method A, quantifying cell–cell distances using centroids, a cell's nearest neighbor may be incorrectly identified because centroids do not capture irregularly shaped cells. Method B, quantifying cell–cell distance by measuring between cell edges, would correctly identify which cell is a given cell's nearest neighbor. At the bottom of (A) is an example where a cell may have as many as four neighbors directly touching it, but looking for the closest centroid would arbitrarily rank equivalent cells in terms of their distance. **(B)** A histogram of how many directly touching neighbors each cell has, peaking at 2-3 per cell. Simply looking for a given cell's "nearest neighbor" would frequently neglect several other equivalently close cells. **(C)** A region of one FOV showing a megakaryocyte (outlined in white) and all its neighbors that are in direct contact (outlined in red). The fact that this megakaryocyte has 15 equivalently close neighbors could not be captured by simply examining cell centroid-to-centroid distances. Scale bar represents 10 microns.

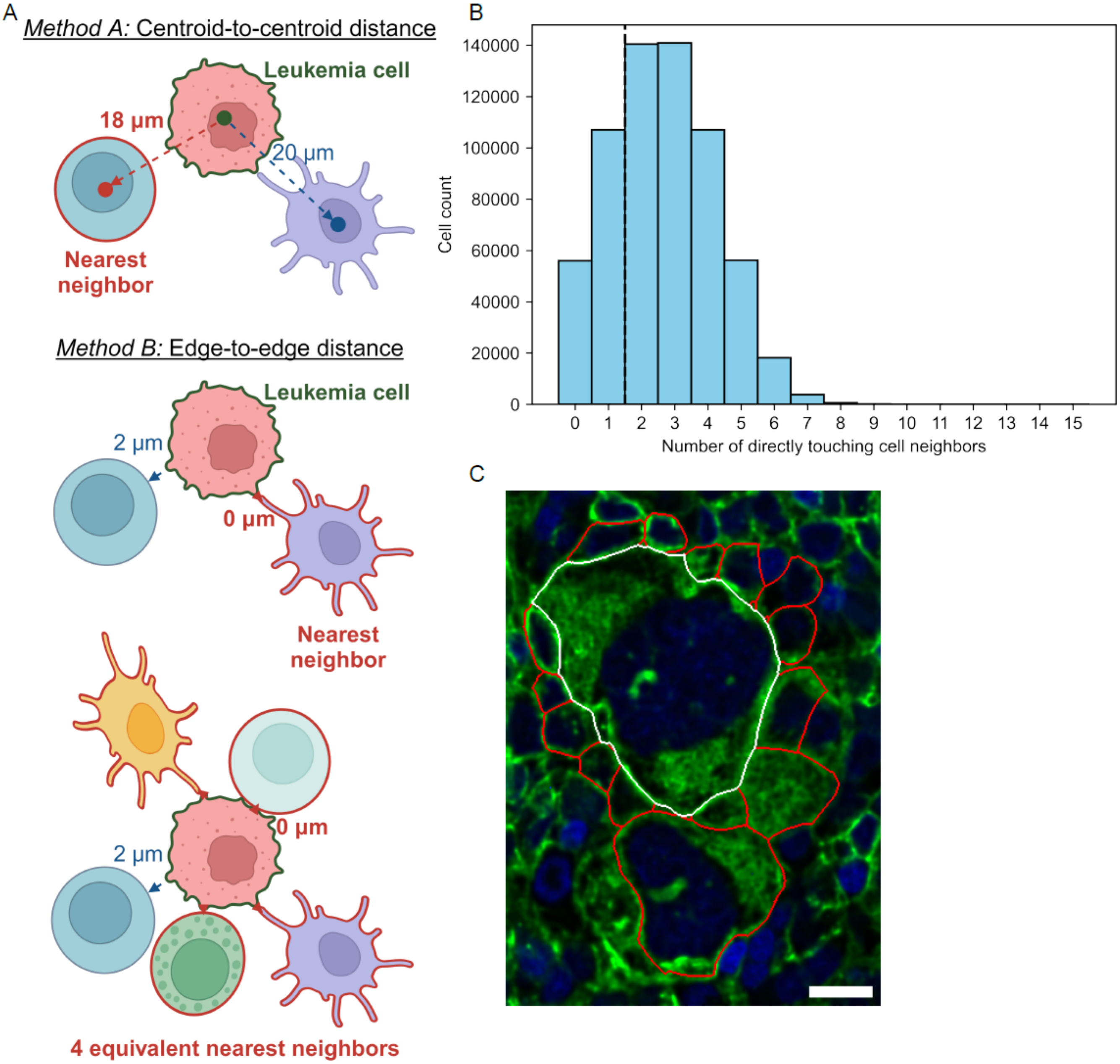

**Figure S9. Linear mixed model baseline comparison. (A)** Comparison of granzyme K+ CD8 effector T cell proportion around leukemia cells at baseline between nonresponders and responders. **(B)** Comparison of $CD14^+$ monocyte proportion around leukemia cells at baseline between nonresponders and responders. **(B)** Comparison of monocyte progenitor cell proportion around leukemia cells at baseline between nonresponders and responders. RR: rate ratio. All p-values in the figure were adjusted p-values considering the size of the model.

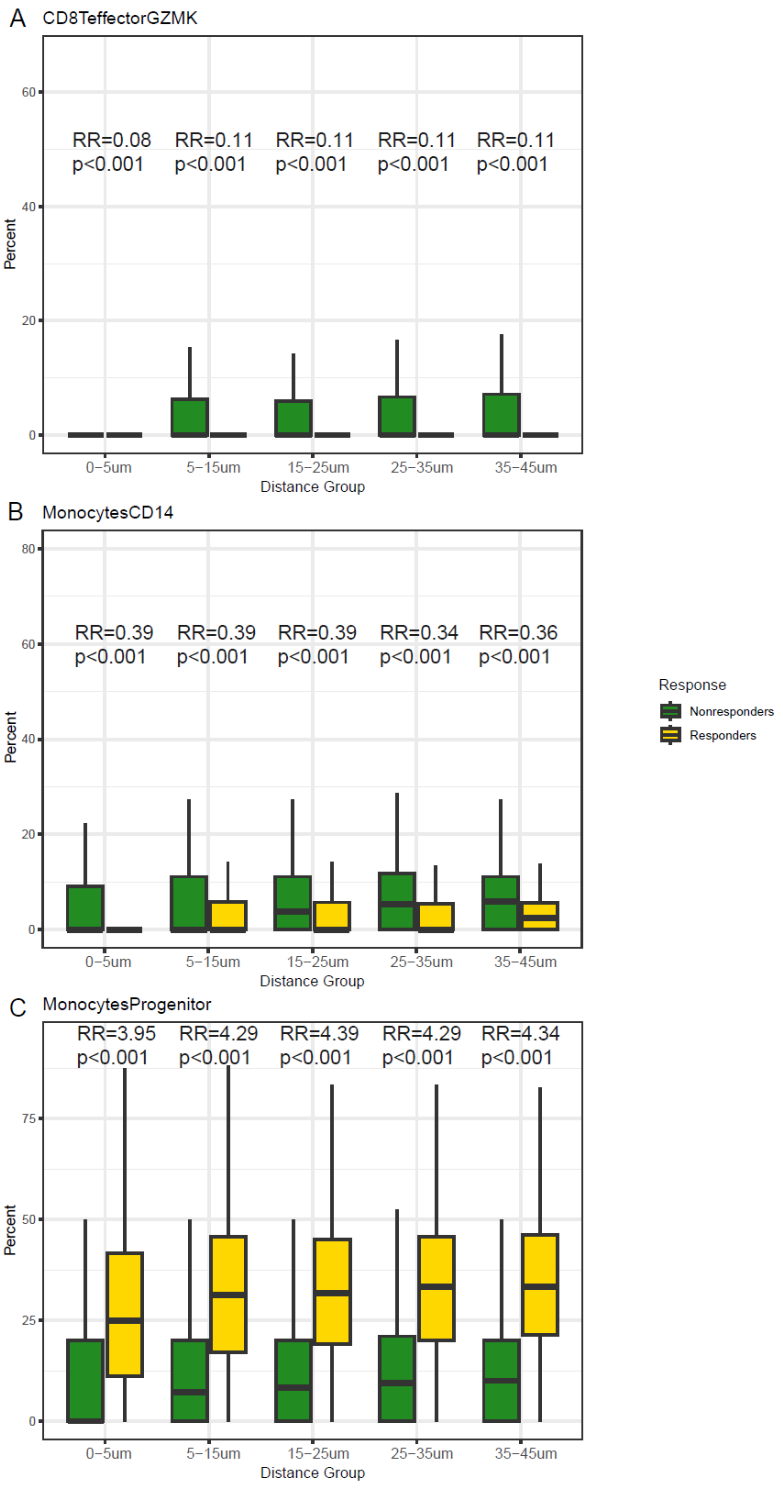

**Figure S10. Schematic depicting cell density shift analysis.** We computed a distribution of distances from a leukemia cell to the nearest of another cell type of interest (T cell in this example) separately at baseline and at the post-ICI time point to create probability density functions (pdfs) for each time point. Subtracting the pdf at baseline from the post-ICI pdf, we obtain the probability density difference across possible leukemia-T cell distances (*x*-axis). This true probability density difference was compared to a null probability density difference created by calculating the distribution of shortest distances from a cell of any type to a T cell. We considered the true probability density difference to be significantly different than expected by random chance if its absolute value was greater than the absolute value of the null probability density difference.

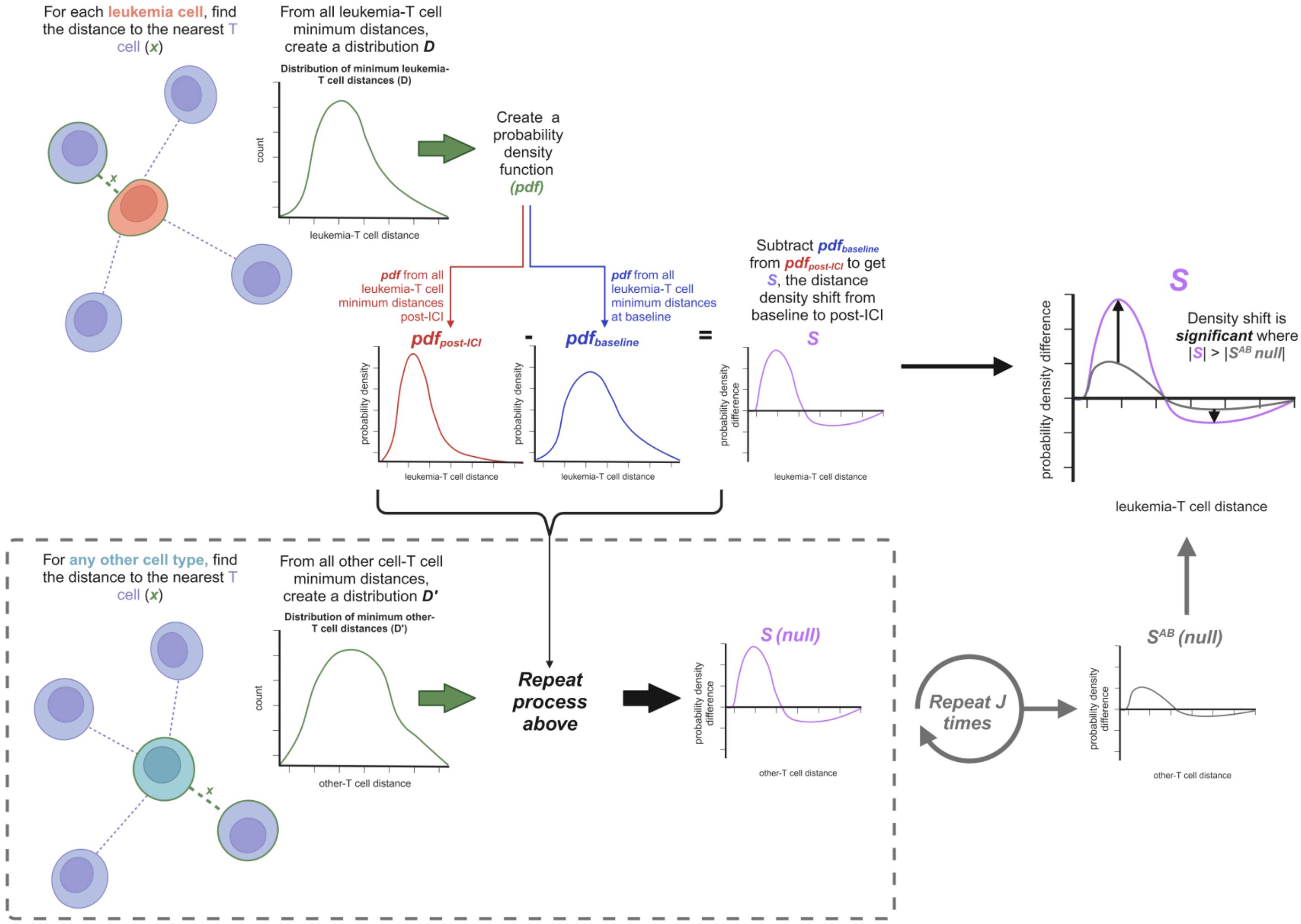

**Figure S11. Cell density shift analysis for each cell type across all patients. (A)** Shows results from all T cell subsets and **(B)** from all other cell types analyzed.

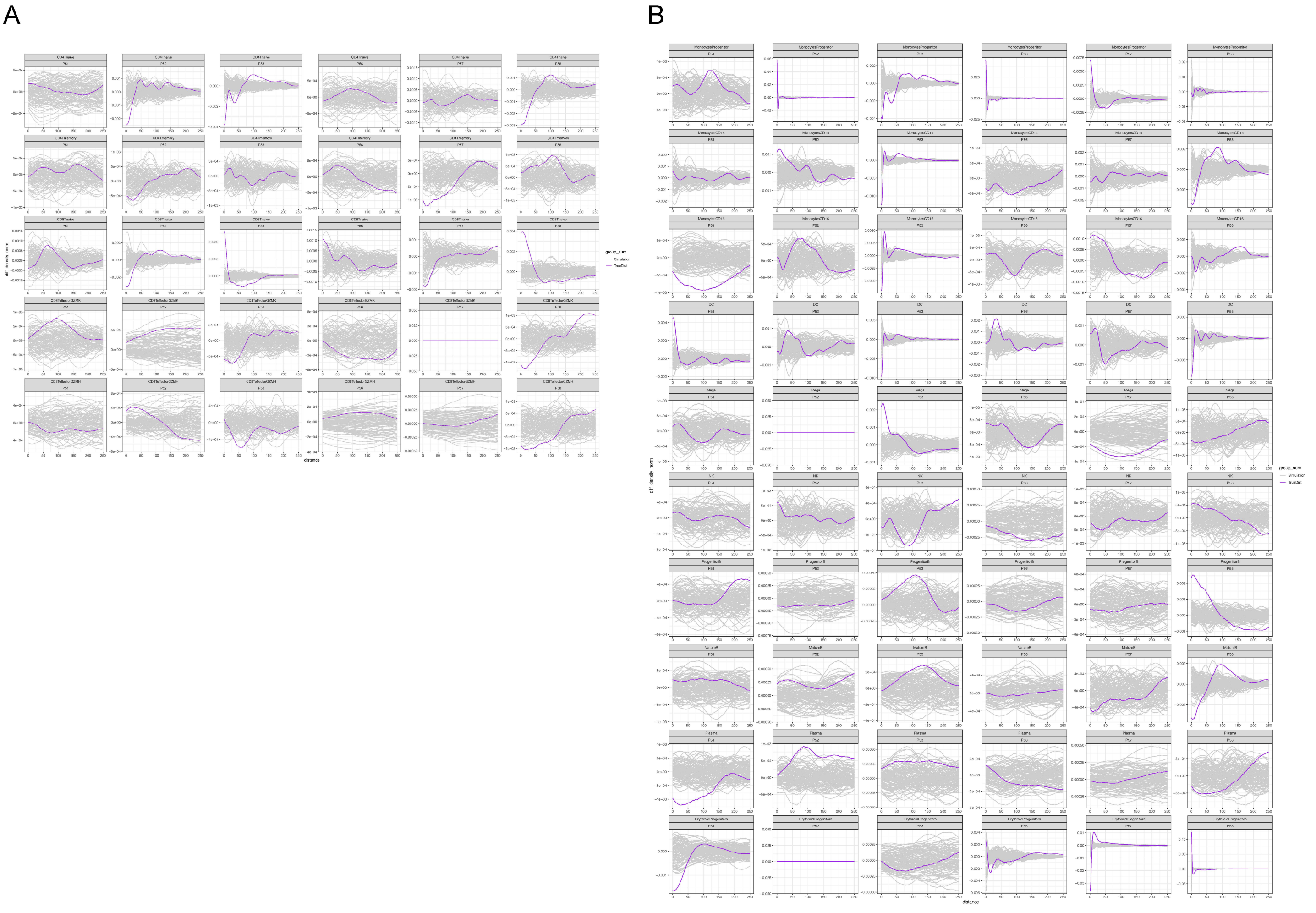
